# Sanctioning of bacterial cheaters by the host plant in nitrogen-fixing symbiosis between *Medicago truncatula* and *Sinorhizobium meliloti*

**DOI:** 10.1101/2024.09.24.614788

**Authors:** Min Chen, Axelle Raisin, Nathalie Judkins, Pierre-Marie Allard, Emmanuel Défossez, Michael Stumpe, Inmaculada Yruela, Manuel Becana, Didier Reinhardt

## Abstract

In plant-microbe interactions, the host plant invests considerable amounts of resources in the microbial partner until the symbiotic machinery is established. If the microbial partner does not reciprocate with a comparable symbiotic benefit, the interaction represents a parasitic relationship. This is thought to elicit a plant’s response to prevent the selective disadvantage of being parasitized by such microbial cheaters. Indeed, negative feedback against bad mutualists, known as sanctioning, has been observed in interactions such as the arbuscular mycorrhizal and legume-rhizobium symbioses. Here, to study sanctioning by the plant host, we manipulate the exchange of resources between the model legume *Medicago truncatula* and its bacterial partner *Sinorhizobium meliloti* by three ways: mutating the bacterial nitrogenase enzyme, replacing nitrogen in the atmosphere with argon gas, and supplying high nitrate to the host. Then, we follow the consequences for the interaction by examining the metabolome, proteome, and phosphoproteome of nodules. We find that such cheating conditions result in sanctioning of the bacterial partner, and observe characteristic shifts including induced defense markers, repressed symbiotic markers, and changes in central metabolism that may be relevant for microbial fitness and that could therefore contribute to sanctioning.

## Introduction

Endosymbiosis with microbial partners that confer a metabolic advantage is extraordinarily successful in plants, with the founding event being the domestication of photosynthetic cyanobacteria that became permanent cellular constituents, the chloroplasts (plastids in non-fixing tissues) (Zimorski et al., 2014). Yet, the majority of plants entertain an endosymbiotic relationship with fungi (arbuscular mycorrhizal fungi, AMF) that provides their host with various essential mineral nutrients (Smith and Read, 2008). Some lineages of the angiosperms (Fabales, Fagales, Cucurbitales, Rosales), have evolved a symbiosis with nitrogen-fixing soil bacteria (rhizobia), in which the bacteria supply the host with ammonia in exchange for fixed carbon in the form of dicarboxylic acids. The best example of this kind is the legume-rhizobium symbiosis (Oldroyd et al., 2011). This symbiotic interaction clearly provides a critical advantage to both partners; however, the dynamics of the interaction are very asymmetrical. While the host plant evolves relatively slowly as an annual or perennial, the bacterial population expands from a single cell per nodule to tens of millions of bacterial cells (bacteroids) within a few weeks (Quides et al., 2021). The resulting conflict between the symbiotic partners has resulted in the evolution of feedback mechanisms that allow the host to control the bacterial community in the nodules (Sachs et al., 2018.

Host risks to become exploited by symbionts with reduced symbiotic efficiency (cheaters). Indeed, theoretical considerations suggest that such symbioses should be inherently unstable, unless the host can enforce symbiotic service from the microbe. The risk of degeneration of symbiotic relationships is particularly high when the microbial partner exhibits very dynamic population dynamics. For example, rhizobial infections with single cells gives rise to nodules with many millions of inhabiting bacteroids, raising the possibility that spontaneous mutations that render bacteria ineffective nitrogen-fixers may confer them a selective advantage resulting from a reduced metabolic burden from the extremely expensive nitrogen fixation process. The resulting risk is two-fold: the individual plant may suffer from a disadvantage compared to conspecific individuals of the same population with better symbionts; and the local population as a whole may suffer over the years from expanding populations of ineffective or even parasitic rhizobia in a local area. Hence, it has been predicted that for the host species, detection of cheaters and their inhibition (or the specific promotion of good mutualists), is of primordial importance for long-term survival of the species (Kiers and Denison, 2008). Indeed, several studies have shown that legume hosts can distinguish rhizobia according to their symbiotic service, and selectively promote good mutualists; however, the implicated mechanisms remain elusive

Sanctioning requires that the host plant can recognize cheaters and that it can exert a negative feedback onto them. Hence, two major questions arise: (1) what are the metabolic consequences of cheating for the host, and which molecule(s) may serve as signals for cheating? and (2) which mechanisms are activated in the host plant to sanction cheaters? The first answer may involve intermediates or end products of N assimilation. The second answer can involve direct inhibition of bacterial growth and cell division, reduced allocation of C (or other essential nutritional requirements), or induction of a defense response involving antibacterial proteins and metabolites.

Here we use a metabolomic and proteomic approach to characterize the interaction between *Medicago truncatula* and *Sinorhizobium meliloti* under conditions that prevent N-fixation (inoculation with non-fixing mutant rhizobia or under N-free atmosphere), and under conditions that suppress symbiosis (high N-supply). With this approach we test hypotheses that could explain the selective repression of cheating rhizobia by the host plant. In particular, we ask whether the plant mounts a defense response, increases the expression of antimicrobial NCR peptides, or alters oxygen and/or nutrient supply to the bacteroids.

To control N fixation in *M. truncatula* nodules, we inoculated plants with either wild-type *S. meliloti* (N-fixing), or with *S. meliloti nifH* mutants that lack an essential component of nitrogenase due to a transposon insertion (non-fixing). In addition, we employed an interaction in which plants were inoculated with wild-type (wt) rhizobia and subsequently cultured in N-free argon atmosphere (non-fixing). In the latter two cases (*nifH*, argon), the lack of N-fixation resulted in altered nodulation. Nodules were smaller, roundish, and with little pink color of leghemoglobin (Lb) (Fig. 1). The smaller and more roundish shape of nodules indicates that they lost indeterminacy, which would normally cause nodules to continuously grow and elongate (Lotocka et al., 2012). While the non-fixing nodules where smaller, they were more abundant and often clustered (Fig. 2), indicating that the plants exhibited an increased tendency to form nodules. The reduced nodule size is probably the result of a negative feedback mechanism that inhibits nodule growth, an effect that we further refer to as sanctioning. The bacteria inhabiting non-fixing nodules are classified as cheaters because they consume resources of the host without reciproking with fixed N. Hence, *nifH* mutants and wt rhizobia in plants under argon atmosphere are both considered cheaters. Increased nodule number under non-fixing conditions may indicate that the lack of N fixation triggers a compensatory mechanism that increases the tendency to become infected by rhizobia.

**Fig. 1.**
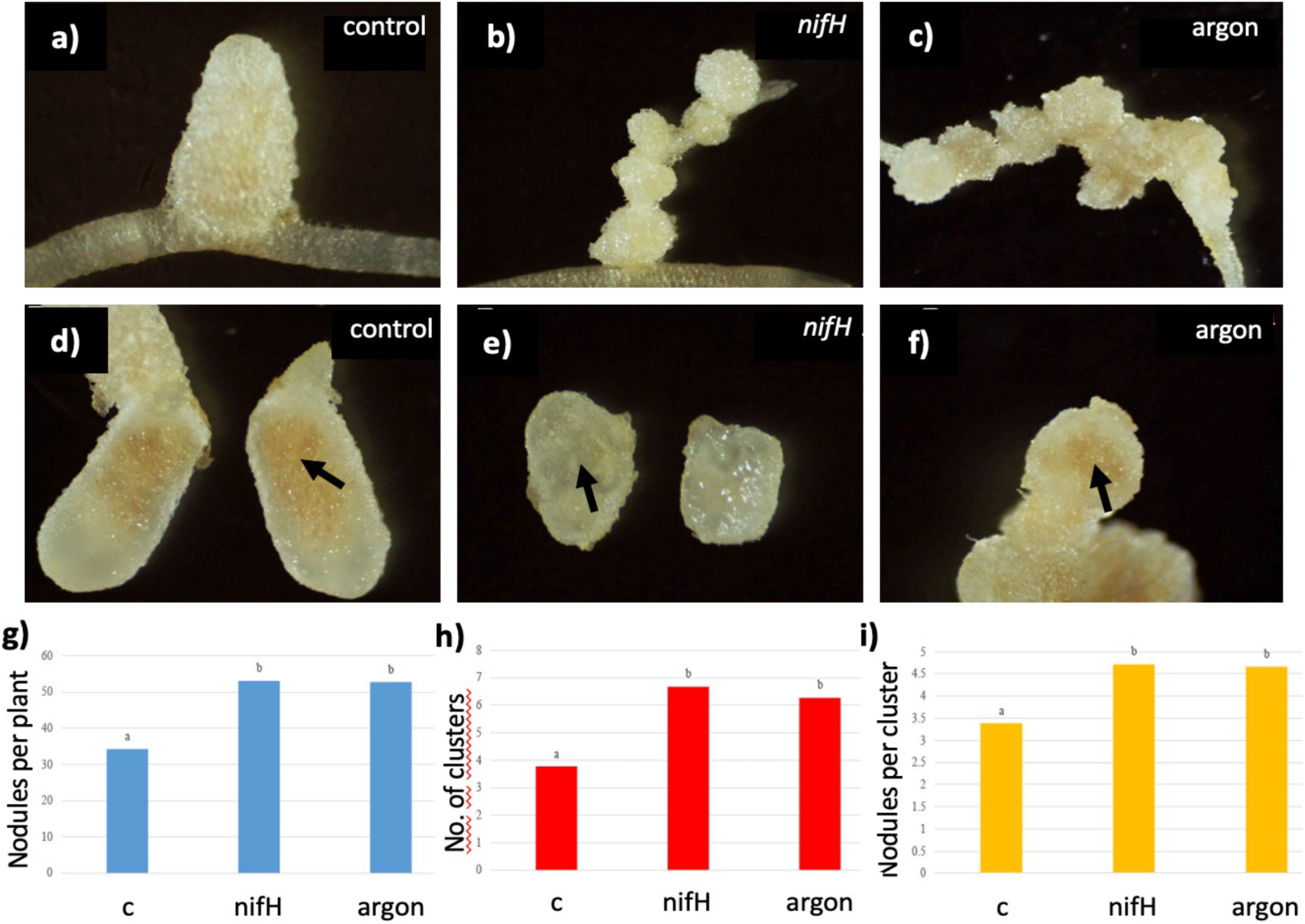
Sanctioning phenomenon against non-fixing rhizobia. **(a,d)** *M. truncatula* nodules inhabited by wt *S. meliloti* and cultured under control conditions. **(b,e)** Nodules inhabited by *nifH* mutant bacteria and cultured under control conditions. **(c,f)** Nodules inhabited by wt *S. meliloti* and cultured under argon atmosphere. **(g-i)** Quantification of total nodule number per plants, number of nodule clusters (groups of neighbouring nodules in direct physical contact, and number of nodules per cluster.

**Fig. 2.**
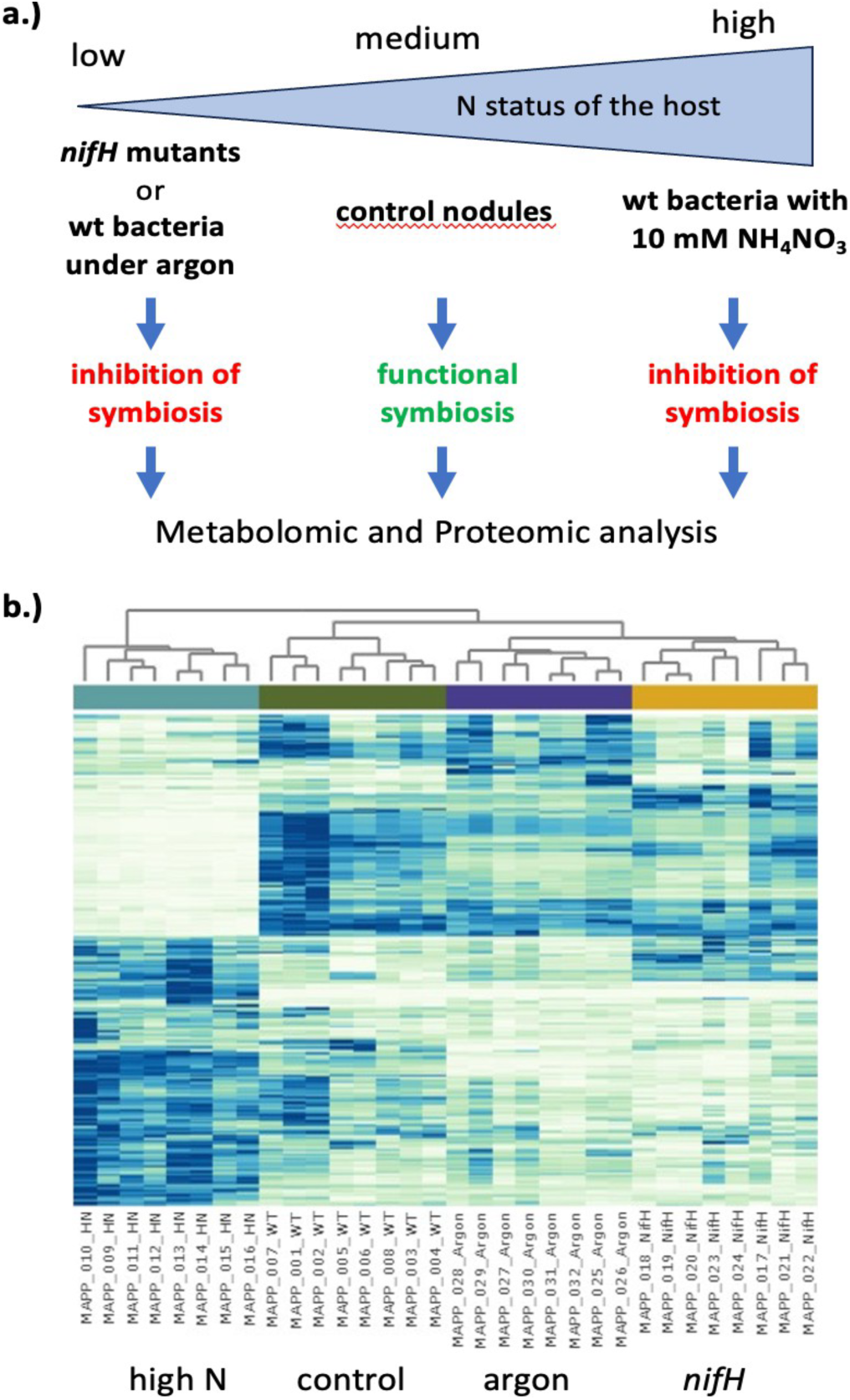
Experimental setting. **(a)** Sanctioning of the bacterial partner by the host is controlled via manipulation of N status. The symbiotic system is either deprived of N (inoculation with *nifH* mutants or exposure to argon atmosphere) or treated with HN (10 mM NH4NO3). In all three cases, symbiosis is aborted compared to control conditions. Nodules harvested from the four growth conditions were harvested for metabolomic and proteomic analyses. **(b)** Global clustering of metabolomic dataset reveals consistent abundance patterns of chemical features. All samples clustered according to their group. A strong shift in abundance is particularly obvious for the HN treatment.

## Results

### Sanctioning: negative feedback regulation of ineffective nodules by the host

Host plants can exert negative feedback regulation onto nodule development when the interaction is not beneficial to the host, for example when rhizobia do not provide fixed N (**Fig. 1**). When *M. truncatula* is inoculated with a *S. meliloti nifH* mutant incapable of N fixation (**Fig. 1b,e**, compare with **Fig. 1a,d**), or if plants inoculated with wt rhizobia are cultured in an N-free argon atmosphere (**Fig. 1c,f**), nodules are small and contain reduced levels of Lb, which controls O_2_ supply to the bacteroids (**Fig. 1d-f**) (Larrainzar et al., 2020). In addition, plants produce excess of these small nodules (**Fig. 1g**), which often occur in dense clusters (**Fig. 1b,c** and **h,i**).

### Metabolomic analysis of sanctioning

To gain insight into the mechanisms of sanctioning, we first performed a metabolomic analysis of nodules under the experimental conditions mentioned above (Fig. 2a). In addition, we included a third condition, high exogenous N supply, that results in inhibition and abortion of nodule symbiosis (Oldroyd and Leyser, 2020). To make the nodules as comparable as possible, we grew all plants (inoculated with wt *S. meliloti* or with a Tn5-tagged *nifH* mutant) together under the same conditions for two weeks. Then, a third of the plants inoculated with wt bacteria were subjected to either argon atmosphere or supplied with 10 mM NH_4_NO_3_ (high N), respectively, whereas one-third of the wt and the *nifH*-inoculated plants continued to grow under control conditions (air, basal nutrient solution).

The global metabolomic analysis of four independent biological replicates, each with *c.* 100 pooled nodules and with two technical replicates per biological replicate, revealed a highly consistent clustering of the data, showing that the dataset is robust and reliable, and that each treatment results in specific metabolomic patterns (**Fig. 2b**). Taking as a proxy the *c.* 10% contribution of bacterial reads in the total transcriptome of nodules (Roux et al., 2014), we assume that *c.* 90% of the metabolites analyzed in our experiments are of plant origin; hence, most of the data will reflect plant metabolites.

To identify molecular features related to specific nutritional N-status and to biological states in nodulation, we compared treatments pairwise and in groups. This allows to identify common patterns of metabolic response under conditions of cheating by non-fixing rhizobia (*nifH* or argon *vs.* controls) from more general mechanisms involved in inhibition of nodule growth (*nifH* or argon or HN *vs.* controls). Furthermore, the HN treatments can serve as a positive control for the metabolic consequences of rich N-supply (HN *vs.* controls) in contrast to N-limitation (*nifH* or argon *vs.* control). Ultimately, this approach allows to separate the metabolic consequences of N-depletion from the mechanisms involved in sanctioning.

### N-related metabolic patterns in response to various N-states

First, the most-discriminant metabolites were determined in volcano plots between the sets of control nodules, i.e. inoculation with wt bacteria in air vs. non-fixing nodules, i.e. inoculation with wt bacteria in argon, or with *nifH* mutants (Fig. 3a). In order to focus on N-fixation-related effects, only metabolites behaving similarly in both argon and *nifH* samples were considered based on automatic annotation of chemical features.

**Fig. 3.**
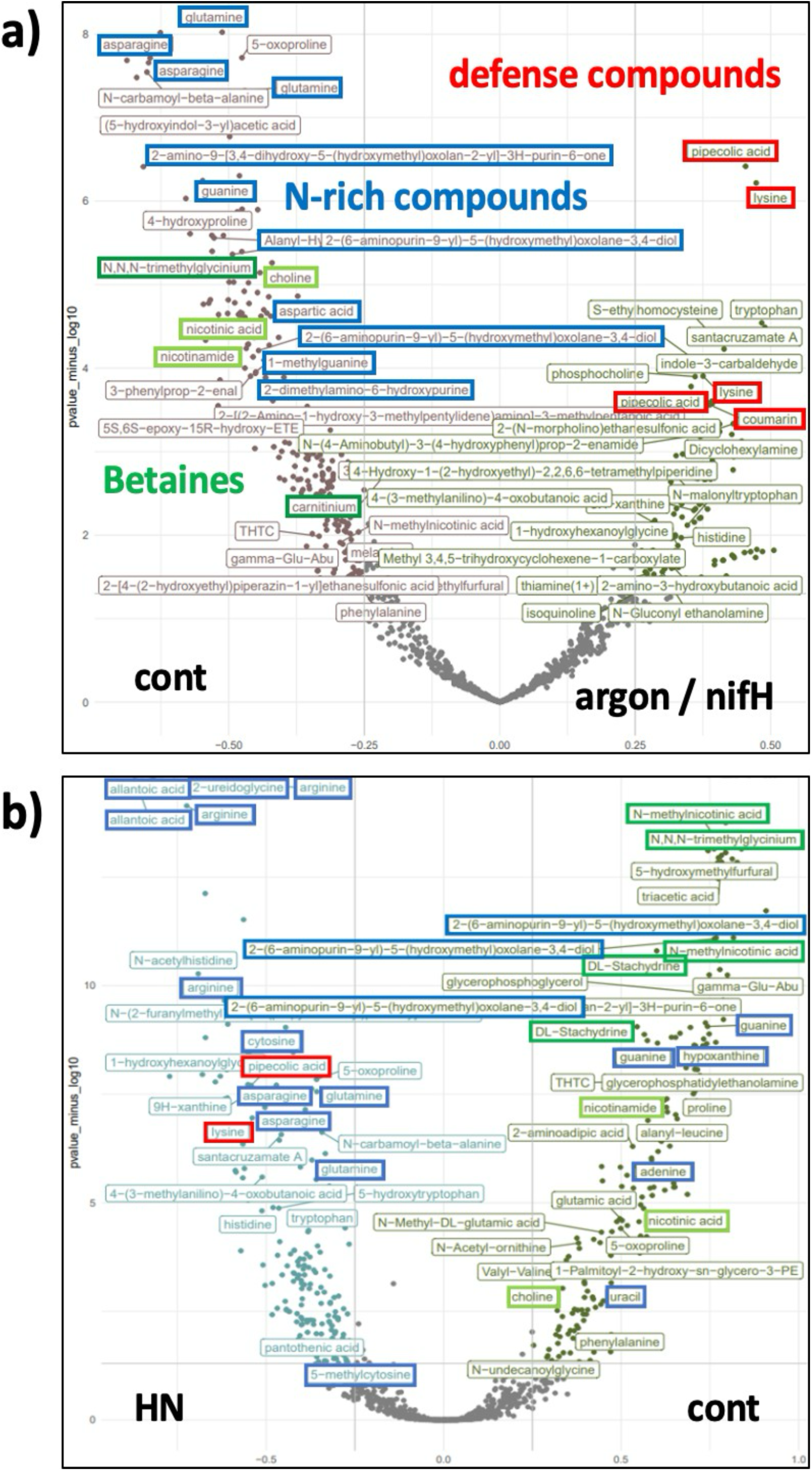
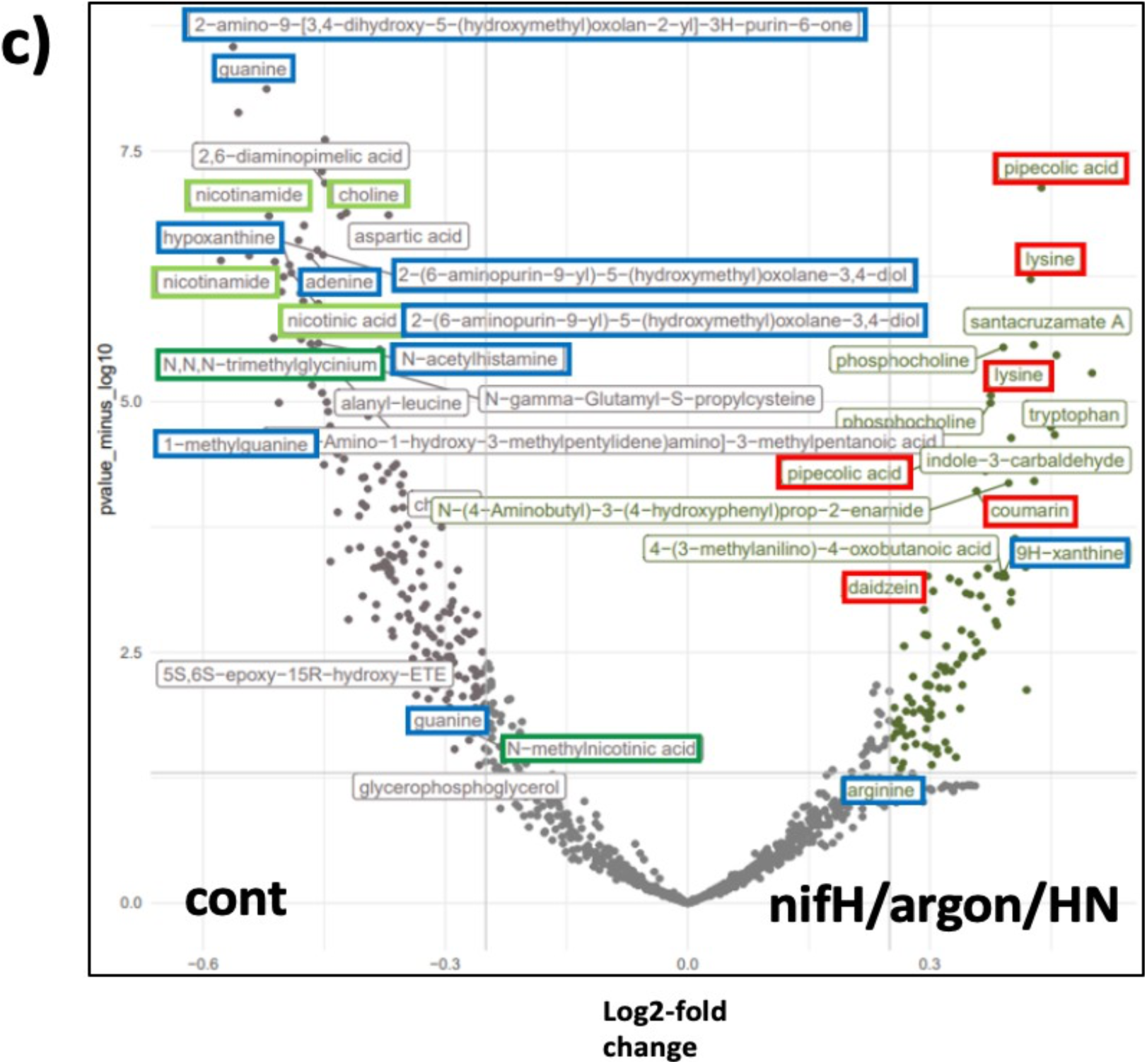
Global assessment of most-discriminating chemical features. **(a)** Volcano plot analysis of all chemical features according to the -log2 -fold change (*x*-axis) and the -log10 *p*-value for the significance of the change (*y*-axis) in the pairing of control nodules (cont) *vs.* both conditions with non-fixing nodules (*nifH* and argon). **(b)** Volcano plot analysis as in **(a)** with the pairing of control (cont) *vs*. high N (HN). **(c)** Volcano plot analysis as in **(a)** comparing control nodules (cont) with all three sanctioning conditions (*nifH*, argon, HN). Note that in **(a)** and **(c)**, only features are shown that were significantly affected both by *nifH* and argon **(a)**, and by all three conditions (*nifH*, argon, HN) **(c)**, respectively.

Consistent with their assumed N-status (**Fig. 2a**), wt samples showed several N-rich intermediates, such as Gln, Asn, and guanine, besides several other purine-related derivatives such as methylguanine (**Fig. 3a**). Sometimes, multiple metabolite features were annotated with the same name, indicative of isomers that have the same mass, but distinct separation kinetics in the HPLC column. In these cases, chemical standards were used to identify the compounds with confidence (see below).

In the comparison of control nodules to those from HN plants, two groups of molecules were particularly discriminative: (i) in HN-samples, several N-rich compounds were observed, and (ii) in control nodules, several molecules were detected with quaternary N atoms, also known as betaines (McNeil et al., 1999). Additional betaines induced in control nodules were stachydrine (N-dimethylproline) and trigonelline (N-methylnicotinic acid), together with their respective precursors Pro, choline, and nicotinic acid (**Fig. 3b**). A related compound found in wt nodules was carnitine (**Fig. 3a**). Betaines and carnitine generally accumulate in response to osmotic stress and may serve as osmoprotectants (Vriezen et al., 2007). Stachydrine is essential for nodulation because bacterial *stc* mutants that cannot catabolize stachydrine are symbiosis-defective (Goldmann et al., 1994).

In comparison to control nodules, the nodules from HN plants accumulated even larger amounts of the amides Gln and Asn, in addition to other N-rich compounds such as Arg, allantoic acid, xanthine, and 2-ureidoglycine (**Fig. 2b**; **Fig. 3a,b**). Ureides such as allantoin and allantoic acid represent the predominant transport form of N in legumes from tropical areas such as soybean (Todd et al., 2006), whereas *M. truncatula* is considered an ’amidic’ species that transports most of its fixed N in the form of Asn (Fischinger and Schulze, 2010). Considering allantoin biosynthesis under high N-supply, two interpretations are possible: it could represent anabolism of a highly efficient storage and transport form of N (with a N:C ratio of 4:4 per molecule) (Thomas and Schrader, 1981), or catabolism of purines, which could be regarded as a sign of senescence (Zrenner et al., 2006; Werner and Witte, 2011). Interestingly, control nodules accumulated guanine (**Fig. 2a,b**), whereas HN nodules accumulated the allantoin precursors xanthine and hypoxanthine, indicative of an N-dependent transition in purine metabolism (see below).

Taken together, this initial metabolomic analysis confirms the assumption that our treatments cover a range of different N-states: N limitation (*nifH*, argon), high nitrate (HN), and an intermediate N status (N-fixing control) plants. Quantitative determination of the primary products of N assimilation Gln and Asn (Xu et al., 2012) showed that controls contained intermediate amounts (resulting from basic nutrient solution with 0.5 mM KNO_3_ and N-fixation), whereas the amide contents were low in *nifH* and argon samples (depending on only exogenous 0.5 mM KNO_3_) and but were consistently augmented in the HN (10 mM NH_4_NO_3_) treatment (**Fig. 4a,b**).

**Fig. 4.**
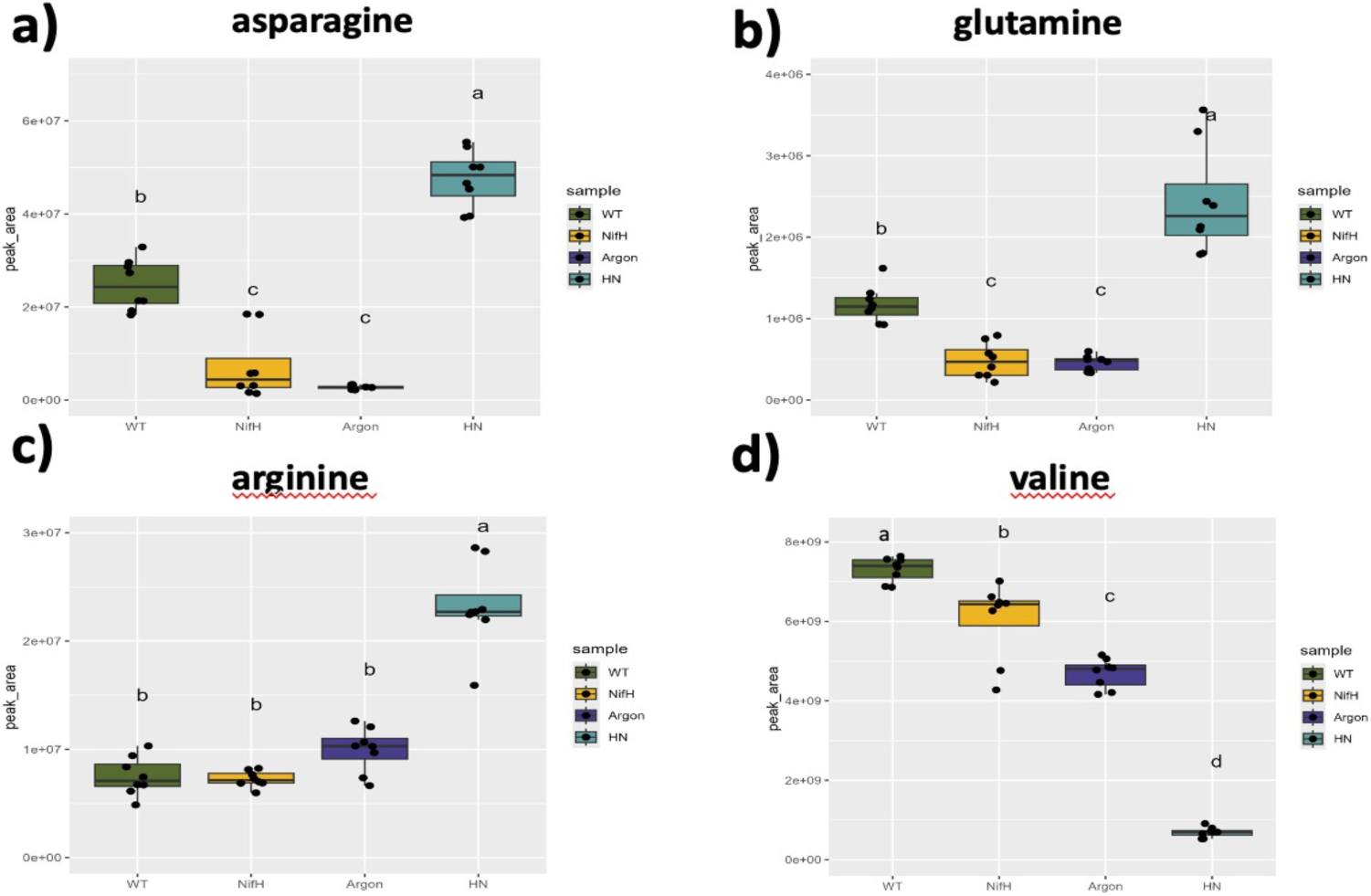
Relative levels of diagnostic amino acids. **(a,b)** The central metabolic intermediates in N-assimilation, Asn and Gln, showed low levels in non-fixing nodules (*nifH* and argon, respectively), high levels in HN and intermediate levels in controls (wt). **(c)** Arg, an amino acid with a high relative N-content (6 C, 4 N), was highest in HN nodules and low in all the others. **(d)** Val, a representative of the three branched amino acids (Leu, Iso, Val), showed highest levels in controls nodules, lowest levels in HN nodules, and intermediate levels in non-fixing nodules (*nifH*, argon).

### Global metabolic signature of sanctioning in nodules

To obtain insight into the pathways involved in sanctioning, we searched for marker metabolites that are significantly affected by argon and *nifH* treatments relative to control nodules. Apart from the strong reduction in N-rich compounds under non-fixing conditions (see above), the most discriminant compounds in argon and *nifH* treatments were the defense signal pipecolic acid and its precursor Lys, which were both strongly induced in sanctioned nodules (**Fig. 3a**). Pipecolate is an established inducer of systemic acquired resistance (SAR) (Návarová et al., 2012; Bernsdorff et al., 2016), which mediates plant immunity against a wide range of pathogens through its mobile derivative *N*-hydroxypipecolic acid (NHP) (Hartmann and Zeier, 2019; Vlot et al., 2021). Interestingly, pipecolate was also a distinguishing feature in the reaction to high N levels (**Fig. 3b**), indicating that defense may become induced in all conditions that are associated with an abortion of symbiosis. Indeed, differential volcano plot analysis of control conditions *vs.* all three conditions with aborted nodules (in a clustered setting) identified pipecolate among the most discriminating factor, together with defense compounds such as coumarin and daidzein (**Fig. 3c**). In contrast, the central regulator of SAR in the shoot, salicylic acid, was not found to be differentially regulated in any of the conditions, in agreement with the finding that defense reactions in the root system are different from the corresponding events in the shoot (Millet et al., 2010; Chuberre et al., 2018; Chen et al., 2021).

### Quantitative assessment of diagnostic pathways potentially involved in sanctioning

Amino acids (aa) play a central role in nodulation due to their role in N-assimilation and due to the dependency of bacteroids on various aa from the host (Prell and Poole, 2006; Prell et al., 2009). As expected from volcano plot analysis (**Fig. 3**), Asn and Gln as primary N-acceptors in N-assimilation, were low in argon and *nifH* samples, and high in NH treatments (**Fig. 4a,b**). The N-rich aa Arg (with 4 N atoms per molecule) was strongly increased in the HN treatment relative to all others (**Fig. 4c**). Strikingly, the levels of the branched aa Val was very low in HN nodules (**Fig. 4d**), indicating that plants may sanction bacteroids by withholding essential aa under high N-conditions.

Next, we considered central C intermediates relevant for bacterial nutrition. The disaccharide trehalose, which is produced primarily by the bacterial partner (Domínguez-Ferreras et al., 2009) (Reibach and Streeter, 1983; Salminen and Streeter, 1986), was decreased under all sanctioning conditions, particularly at HN, whereas the effect of argon was not significant (Fig. 5a). A similar trend was observed with the plant-borne disaccharide sucrose (Fig. 5b), and with the carbon currency supplied to the bacteroids succinic acid (Fig. 5c), consistent with a generally decreased supply of C to nodules (sucrose), and to the bacteroids (succinate), resulting in depleted bacterial C resources (trehalose). The defense signal pipecolic acid was significantly increased under all sanctioning conditions (Fig. 5d), together with its precursor Lys (Fig. 5e) (Hartmann and Zeier, 2019). Similarly, all treatments showed increased levels of the defense compound coumarin (Fig. 5f).

**Fig. 5.**
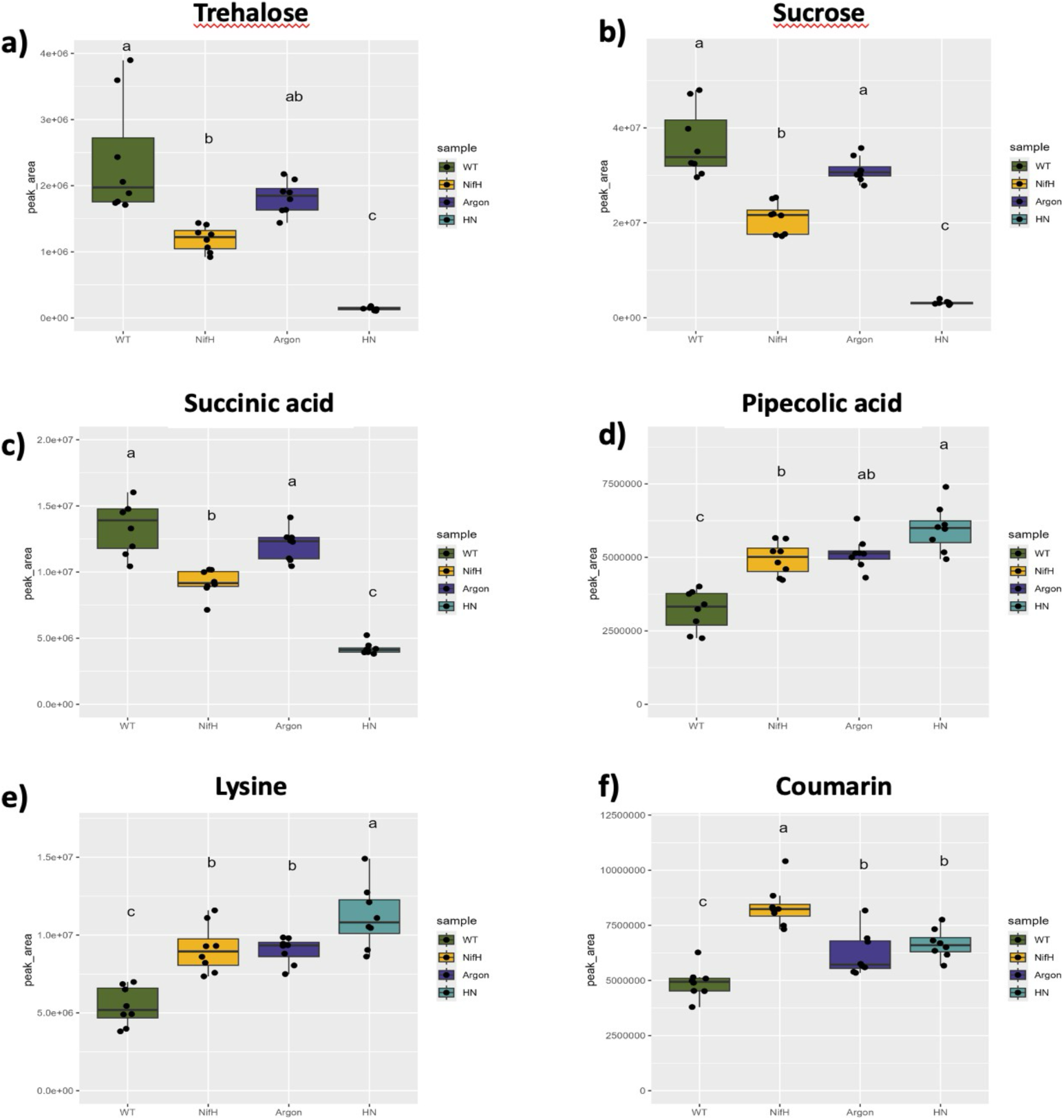
Metabolic markers for carbon resources and defense intermediates. **(a)** Trehalose, a disaccharide produced primarily by the rhizobial partner, decreased in all sanctioning conditions, although not significantly in argon. **(b)** Sucrose, a crucial disaccharide coming from the host, showed a parallel pattern to trehalose (compare **a** and **b**). **(c)** Succinic acid, a C source for the bacteroids, showed a parallel pattern to trehalose (compare **a** and c). **(d-f)** The defense signal pipecolic acid, its precursor Lys, and the defense compound coumarin were induced in all sanctioning conditions.

Quantitative assessment of purines and their catabolites revealed high levels of hypoxanthine and guanine in control nodules, whereas they were decreased in all other treatments (**Fig. 6a,b**). The purine catabolites allantoin and allantoic acid, which are normally not detected in nodules of *M. truncatula*, were strongly induced under HN conditions (**Fig. 6c,d**). This indicates a transition from purine anabolism under control conditions to purine degradation at HN (**Fig. S1**). Alternatively, high N supply may induce a new pathway for N-transport related to the ureide pathway known from tropical legumes such as soybean (Todd et al., 2006).

**Fig. 6.**
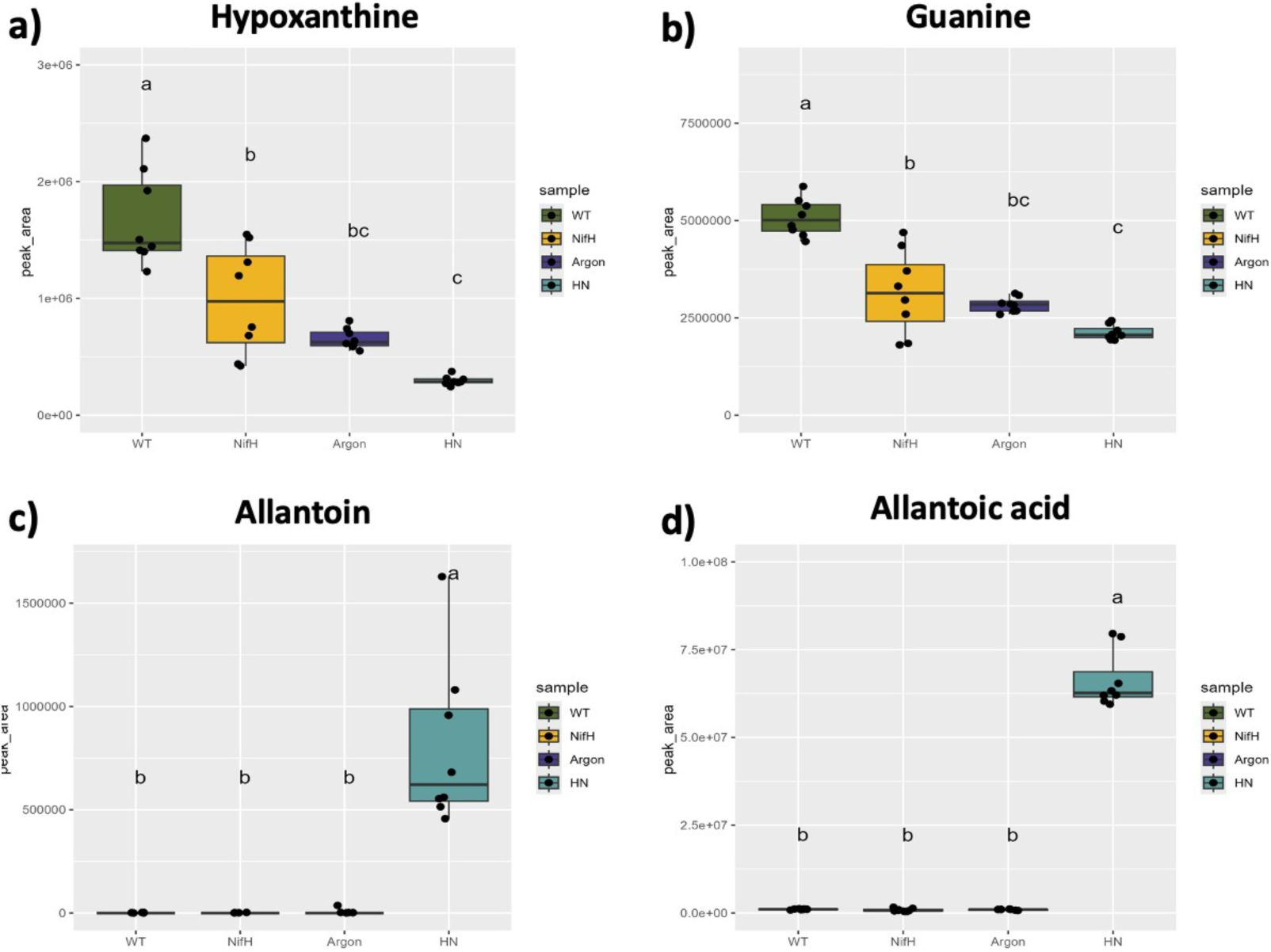
Purines and their catabolites. **(a,b)** The purines hypoxanthine and guanine were increased in control nodules and reduced in all sanctioning conditions (*nifH*, argon, HN). **(c,d)** The purine catabolites allantoin and allantoic acid were detected exclusively under HN conditions.

As for the betaines, there was a strong of glycine betaine (**Fig. 7b**), stachydrine (**Fig. 7d**), and trigonelline (**Fig. 7f**) under HN conditions, whereas argon and *nifH* nodules were similar to the controls (**Fig. 7d,f**). The respective precursors choline (**Fig. 7a**), Pro (**Fig. 7c**), and nicotinic acid (niacin) (**Fig. 7e**) showed similar trends, in particular in the HN treatment. Taken together, betaine levels are predominantly decreased under conditions of high N supply. The related compound carnitine exhibited a moderate trend to decrease under sanctioning (**Fig. 7g**).

**Fig. 7.**
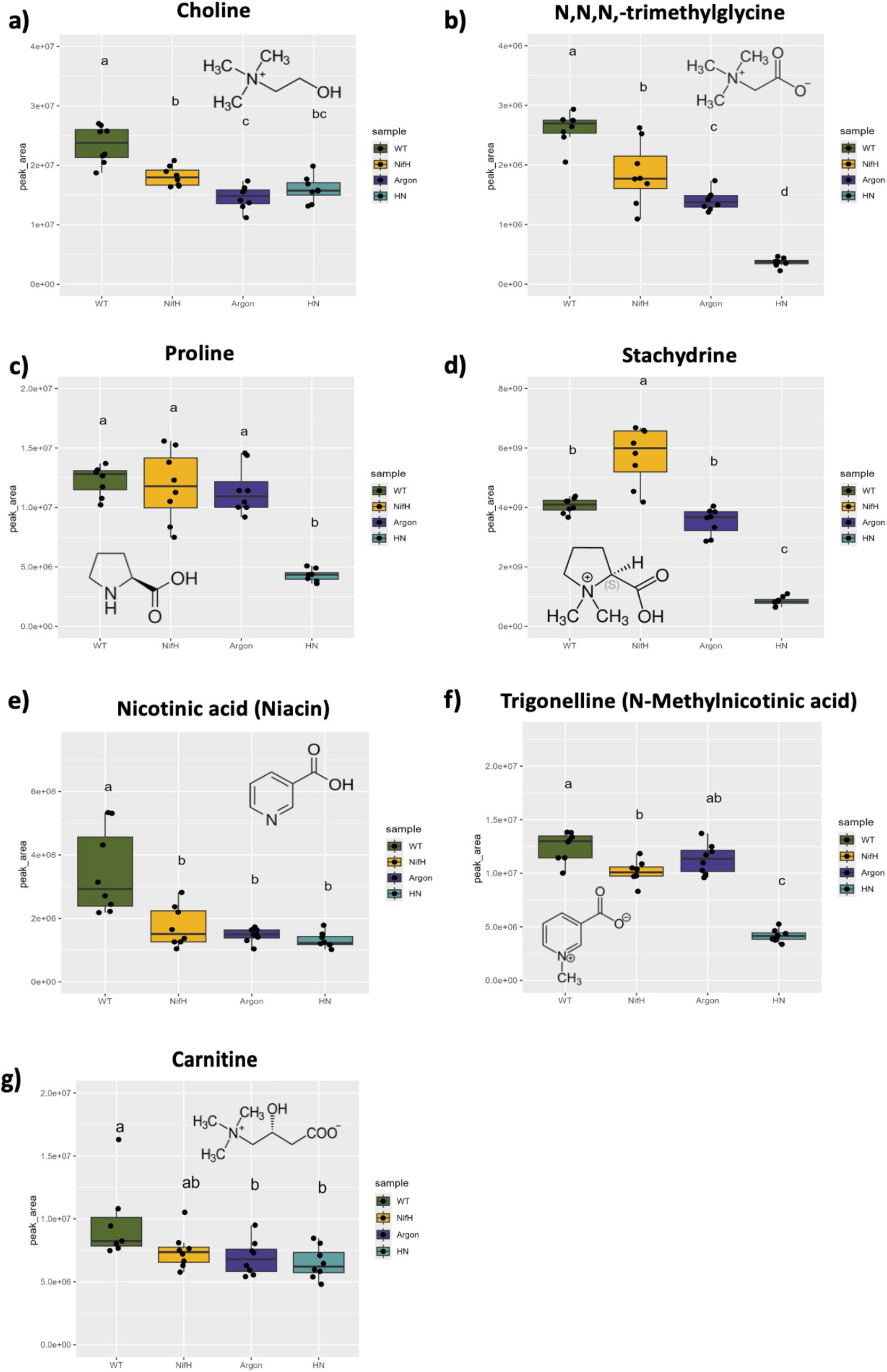
Betaines and their precursors. Glycinebetaine (N,N,N-trimethylglycine) and its precursor choline, as well as nicotinic acid (niacin), were reduced in all three sanctioning conditions (*nifH*, argon, HN). Stachydrine and its precursor Pro were reduced in HN conditions; trigonelline was reduced in *nifH* and HN conditions; and carnitin, which has a structure similar to glycinebetaine, was reduced in argon and HN conditions.

### Proteomic analysis of sanctioning

Metabolomic analysis under sanctioning conditions revealed major shifts in several groups of metabolites, which are likely to reflect reprogramming of plant metabolism (purine and its derivatives, betaines, defense compounds carbohydrates etc.). To obtain a deeper insight into the changes triggered during sanctioning, we explored the proteomic space of the two partners to search for signatures of mechanisms that could be relevant for sanctioning. Based on metabolomic evidence, and inspired by information from the literature (Udvardi and Kahn, 1993; Denison, 2000), we mainly considered three complementary (mutually non-exclusive) hypotheses: (i) sanctioning is associated with nutritional shifts (e.g. lower C supply to rhizobia); (ii) sanctioning is accompanied by repression of symbiotic mechanisms; and (iii) sanctioning involves the induction of a defense response against the rhizobia.

We performed two independent proteomics experiments. In the first one, plants were inoculated with either wt *S. meliloti*, with the non-fixing Tn5-insertion *nifH* mutant of *S. meliloti* (Ruvkun et al., 1982), or under argon atmosphere (see above) for three weeks. In a second experiment, argon was applied after a two-week period of development under control conditions to have similar sample material (comparable number and amount of nodules), and to investigate the more immediate consequences of sanctioning than in a continuous setting of three weeks. In this experiment, we also included a high N (10 mM NH_4_NO_3_) treatment as in the metabolomics experiment described above. In addition, we included free-living bacteria in liquid culture and non-inoculated roots as respective controls for the non-symbiotic status of rhizobia and *M. truncatula*. Global clustering analysis for GO-terms sigificantly affected by both sanctioning treatments (*nifH* and argon) *vs*. control nodules in the 3 wk-treatment (**Fig. S2**), and for GO-terms significantly affected by all three sanctioning treatments (*nifH*, argon, HN) *vs.* control nodules in the 7 day-treatment (**Fig. S3**) revealed that the treatments caused reproducible changes in proteomic patterns.

Analysis of the clusters of GO-terms significantly affected by *nifH* and argon revealed induction of a high proportion of defense-related terms (**Table S1**), whereas the down-regulated GO-terms exhibited several symbiosis-related items (**Table S2**). Mapping induced defense markers on the KEGG map “defense” showed that induction occurred at different levels of defense (**Fig. S4**). To corroborate this proteomic pattern, we performed volcano plot analysis with genes in the GO-term “defense” and with the symbiosis-specific NCR peptides. Comparison of nodules with non-inoculated roots revealed that NCR peptides are mostly nodule-specific (only 1 NCR peptide was expressed also in roots), and that roots exhibit a higher defense status than nodules (**Fig. 8a**), in agreement with the notion that establishment of symbiosis involves a repression of defense pathways (Benezech et al., 2020). In contrast, defense markers were induced in nodules colonized by *nifH*, and NCR peptide expression was reduced (**Fig. 8b**). A very similar pattern was found in nodules grown under argon atmosphere (**Fig. 8c**). Taken together, these results confirm the notion that sanctioning involves increased defense and decreased symbiotic compatibility.

**Fig. 8.**
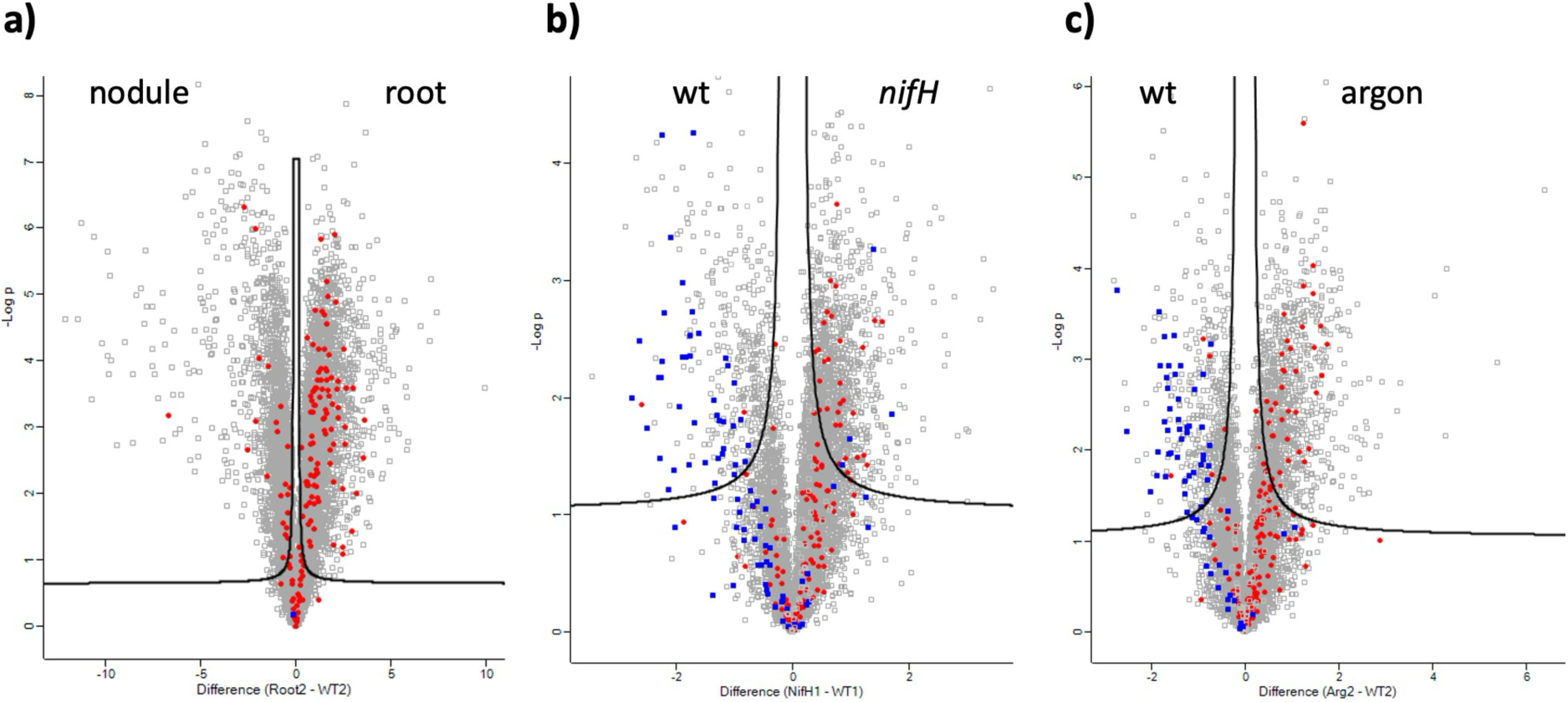
Expression of NCRs and defense-related proteins. Volcano plot analysis showing the differential relative abundances (*x*-axis, difference of -log2; *y*-axis -log10 of *p*-values) of defense-related proteins (red dots) and NCRs (blue dots) in the comparison of **(a)** roots *versus* nodules, **(b)** *nifH* nodules *vs.* wt nodules, and **(c)** argon *vs.* control nodules.

Considering all three sanctioning conditions (including HN), half of the top 10 significantly overrepresented GO-term comprised oxidative pathways including oxidativ stress (**Fig. S5**). A targeted search for homologues of functionally characterized defense markers confirmed that many of them are induced relative to controls under several sanctioning conditions in both experiments (**Table S3**).

A general defense response that is well-characterized at the level of its induction in host plants and at the level of the protective effects against pathogenic microbes is lignin deposition in the cell walls (Lee et al., 2019; Ma, 2024). Lignin biosynthetic enzymes have been characterized in detail (**Fig. S6**) (Vanholme et al., 2019), and the respective genes have been identified in many land plant species, including *M. truncatula* (Ha et al., 2023). Proteomic analysis revealed that several lignin biosynthetic enzymes were induced under sanctioning conditions, although the changes were significant only in the case of PAL, CA4H, 4CL, COMT, CCoAOMT and HCT, (**Table S4**). Analysis of the transcriptional regulation of the respective genes by quantitative real-time RT-PCR (qPCR), revealed an even higher induction of most of the biosynthetic genes (**Table S4**); in particular, of the upstream genes PAL, C4H, 4CL, and C3H, and CCoAOMT that control the flux into the lignin-specific part of the pathway downstream of CCR (Xie et al., 2018; Vanholme et al., 2019).

Among the symbiosis-specific elements in nodulation, the NCR-peptides (NCRs) play a central role. *M. truncatula* encodes several hundreds of NCR genes that are required for the proper development of the bacterial partner for effective symbiosis. Mutations in individual NCRs can result in aborted symbiosis (Horvath et al., 2015; Kim et al., 2015), indicating that they act non-redundantly and fulfill specific functions in bacteroid determination. Among a list of 492 predicted NCR genes, we found 383 to be expressed at the protein level in nodules under control conditions (**Table S5**). Among 154 NCRs that were expressed in nodules under all conditions >100 were significantly repressed in at least 2 of the three sanctioning conditions, whereas only seven were significantly induced in at least two of the sanctioning conditions (**Table S6**).

A second group of host proteins with a central role in symbiosis are Lbs. These hemoproteins regulate O_2_ flux in nodules to reconcile the high rates of bacterial respiration necessary for the energy-demanding nitrogenase reaction with the requirement of low O_2_ partial pressure to avoid oxidation of the iron-sulfur center of the nitrogenase enzyme (Gallon, 1981). *M. truncatula* has 12 Lb genes (Berger et al., 2020; Larrainzar et al., 2020; Jiang et al., 2021), of which 10 were found to be expressed in nodules (**Table S7**). All of these were significantly repressed under sanctioning conditions (**Table S7**), consistent with the reduced appearance of the pink pigment in the nodules (**Fig. 1**), suggesting that either bacterial respiration or nitrogenase activity (or both) must be considerably affected. In addition to reduced Lb protein levels, we found six of them to be phosphorylated at conserved positions (S13, S14, S50, S56, and S92) (**Table S8**; **Fig. S7**). Interestingly, the six genes encoding the phosphorylated Lb proteins clustered together in a promoter sequence comparison (**Fig. S8**) (Jiang et al., 2021), indicating that these genes may have a common evolutionary origin and shared patterns of regulatory elements in their promoters.

### Proteomic signature of a transition in metabolism

Because several betaine-type zwitterion metabolites with quaternary nitrogen atoms were differentially regulated during sanctioning, and since stachydrine is required for functional symbiosis (Goldmann et al., 1994), we interrogated the bacterial and plant proteome for enzymes and transporters related to betaine. Based on the gene complement in the two partners, glycine betaine can potentially be produced by both partners, whereas stachydrine and trigonelline are likely to be of plant origin. Rhizobia have a large number of betaine and choline transporters (Table S8) and have a dedicated locus for catabolism of stachydrine and other betaines (Goldmann et al., 1991; Phillips et al., 1998), thus corroborating the view the betaines are mainly produced by the host and taken up by the rhizobia (Trinchant et al., 2004). We detected peptides of most of the betaine-related genes in *S. meliloti* (Table S9), but only one of them showed a consistent response to the sanctioning conditions in *nifH* and argon nodules (Table S8).

Considering the plant partner, several genes involved in betaine, choline, and carnitine metabolism were detected but only one of them showed a similar response in protein levels under all sanctioning conditions (**Table S10**). These results suggest that betaine metabolism in nodules is not regulated by the abundance of the involved enzymes, but rather at the post-translational level, or, perhaps, by regulation of the substrate concentrations.

### Proteomic signature of a transition in purine metabolism

Purine metabolism is central to bacteria and plants, and both have a large set of enzymes and transporters dedicated to this pathway (**Tables S11 and S12**). *S. meliloti* has a number of allantoin transporters that are expressed in the fixation zone of nodules (Roux et al., 2014). Moreover, legumes are known to produce purine derivatives (ureides) that can function as N-transport form in tropical legumes (Todd et al., 2006). We found little change in the levels of purine- and ureide-related proteins in either organism, indicating that sanctioning does not impinge on these pathways by altering protein levels.

### Proteomic signature of changes in central carbon metabolism

A conceivable sanctioning mechanism against cheater bacteria could be the withdrawal of essential resources such as reduced C or essential aa. Indeed, the levels of central aa and C metabolites were affected by sanctioning (**Figs. 4, 5**). Hence, we searched for proteins that are involved in C metabolism such as enzymes of glycolysis and the tricarboxylic acid cycle. Notably, very little changes were observed in these pathways in both partners (**Tables S13 and S14**), consistent with the commonly observed phenomenon of post-translational regulation of primary C metabolism in plants (Plaxton, 1996; Daloso et al., 2015; Niehaus, 2021). Taken together, the results from proteomic analysis show little evidence for large-scale transitions in enzymes of C metabolism, suggesting that the effects observed at the level of NCRs, Lbs, and lignin-related proteins represent specifically regulated phenomena.

### Phosphoproteome of sanctioning

Because many proteins, including enzymes of central metabolism (e.g. glycolysis), are under control of phosphorylation, we assessed the phosphoproteome of nodules under control conditions and under the three sanctioning conditions (*nifH*, argon, HN). A total of 13,684 tryptic peptides showed differential phosphorylation (Table S15). The 100 top hits (0.7%; sorted according to increased phosphorylation in *nifH* nodules), representing 69 different proteins, featured four main groups (Table S16): The first group is related to defense, including a MAPKKK, calcium and oxidative signaling, and the defense regulator RIN4, which was phosphorylated on a conserved serine residue (Ser85). In Arabidopsis, this residue is phosphorylated by a dedicated immune receptor (RPM1), resulting in increased disease resistance (Liu, 2011).

The second group contained essential elements of nodule functioning, such as Lb (compare with **Table S8**) and an NCR peptide. The third group consisted of elements from central sugar and N metabolism, which regulate C supply to nodules and N assimilation in the host, as well as Pro biosynthesis. The fourth group contained regulators of cellular dynamics, including proteins involved in secretion, membrane transport, membrane dynamics, and cytoskeletal elements. Finally, a conspicuous pair of zinc finger proteins of unknown function was represented by six phosphopeptides that were increased by sanctioning. While the relevance of most of these phosphorylations for protein function is unknown, it shows that central elements of pathways that are influenced by sanctioning are subject to post-translational regulation.

## Discussion

Symbiosis is a very costly arrangement because it can absorb up to 20% of photosynthetic capacity (Jakobsen and Rosendahl, 1990) and consumes large amounts of resources during the establishment of the symbiotic machinery required for sustaining the microbial partner, and for nutrient exchange surfaces and interfaces. It is therefore not surprising that evolution has created mechanisms that prevent symbiosis under conditions under which it does not confer a benefit to the plant, or when it even turns into a parasitic relationship. Under such conditions, plants activate feedback mechanisms that inhibit microbial proliferation and ultimately result in abortion of symbiosis, a phenomenon known as sanctioning of the microbe by the host.

Sanctioning has been observed in many symbiotic associations under various conditions. In the strict sense, the term can be confined to cases where one partner (usually the microbe) provides low levels of symbiotic benefit, which results in a corresponding reduction of the benefit of the other partner (usually the plant). Here, we use a wider definition of sanctioning which includes any situation that results in a negative symbiotic benefit, not only due to reduced service from one partner, but also due to free access of the resources in a symbiosis-independent way (e.g. by exogenous fertilizer supply). Here, we have compared three sanctioning conditions that are all known to result in inhibition of symbiosis, two with reduced N fixation (*nifH*, argon), and one with free access to saturating N fertilization (HN). We then searched for common patterns of sanctioning, and for specific induction of pathways related to the lack of N-fixation and related to saturating N supply.

### Sanctioning via induced defense mechanisms

The most prominently induced metabolic feature shared among all three sanctioning conditions was the defense signal Pip together with its precursor Lys (**Fig. 3c, 5d,e**) and the antimicrobial secondary metabolites coumarin (Stringlis et al., 2019) and daidzein, the precursor of the phytoalexin medicarpin (Yu et al., 2000). Likewise, global GO-term analysis of proteomic results listed defense markers and immune pathways as highly induced in argon and *nifH* nodules (**Table S1; Fig. 8**), and all three sanctioning conditions induced several GO terms involving oxidative stress (**Fig. S5**). Apart from these non-targeted global analyses, induction of defense markers was corroborated by analysis of lignin-related enzymes (**Table S5**), and phosphorylation of defense proteins such as the conserved immune regulator RIN4 (**Table S16**). RIN4 is of particular interest, because it represents a phosphoswitch that can increase, or reduce, immunity of Arabidopsis depending on the site of phosphorylation (Chung et al., 2014; Toruño et al., 2019).

Strikingly, a recent study documented that RIN4 is required for nodule symbiosis in soybean, suggesting that it gates the defense response towards either symbiosis or disease resistance (Toth et al., 2023). In *M. truncatula*, a total of 17 phosphopeptides of RIN4 were detected (**Fig. 9a**), with one phosphopeptide (Ser85, red in **Fig. 9a**) being among the 100 most induced phosphorylation sites (**Table S16**). A dephosphorylated site was detected at Ser140 (blue in **Fig. 9a**), while a symbiosis-related phosphorylation site in a conserved legume-specific motif (GRDSP) (Toth et al., 2023) showed intermediate phosphorylation (green in **Fig. 9b**). RIN4 in soybean (*Glycine max*) interacts with the symbiosis receptors NFR1 and SYMRK and is phoyphorylated by the latter in the GRDSP-motif (Ser143 in soybean; Ser158 in *M. truncatula*) (Toth et al., 2023). It remains to be seen whether secondary phosphorylations observed in our experiments interfere with the symbiosis-promoting phosphorylation site in the GRDSP motif.

**Fig. 9.**
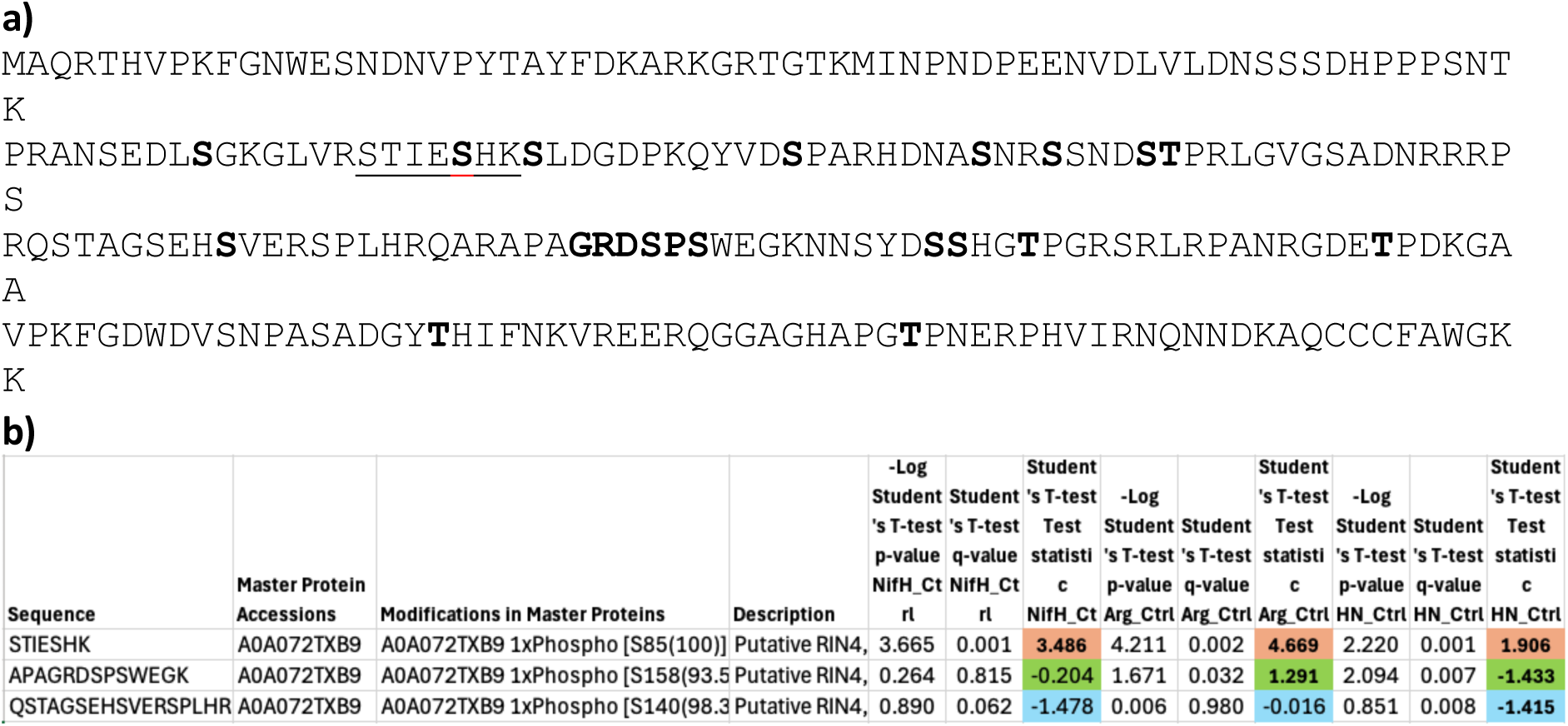
Phosphorylation pattern of *M. truncatula* RIN4. **(a)** All phosphorylation sites are highlighted in color. Induced and reduced phosphorylation are highlighted in red and blue, respectively. The conserved phosphosite in the legume-specific GRDSP motif (Toth et al., 2023) is highlighted in green. **(b)** Quantification of the phosphorylation level of the three phosphosites highlighted in red, green, and blue in **(a).** The relative difference in expression level (difference -log2) is highlighted in the respective colors as in **(a).**

### Sanctioning by repression of symbiotic pathways

Besides the induction of defense, the repression of pathways required for symbiosis was clearly evident for nutritional pathways (sucrose, trehalose, succinic acid) (Fig. 5), as well as for the essential branched amino acid valine (Fig. 4). Furthermore, the general repression of hundreds of NCRs (Table S16) suggests that this phenomenon alone would suffice to abort symbiosis, since even mutations of individual NCRs is sufficient to interfere with nodulation (Horvath et al., 2015; Kim et al., 2015). Finally, reduced Lb levels are likely to interfere with symbiosis since down-regulation or knock-down of Lb results in abortion of symbiosis (Ott et al., 2005; Wang et al., 2019). The significance of Lb phosphorylation is not clear, but the phenomenon has been observed earlier in *M. truncatula* (Marx et al., 2016) and other legumes. Phosphosites S13, S14, S50, and S56 on the surface of the protein might influence interactions with other cellular constituents and molecules, or Lb stability, while phosphosite S92 in the vicinity of the heme coordination center (Fig. S7) may influence O_2_ binding kinetics (affinity, stability).

### A role for betaines in sanctioning?

An unexpected observation was the fact that non-targeted metabolomic analysis identified several zwitterion metabolites with quaternary N-atoms (glycinebetaine, stachydrine, trigonelline, carnitine) as typical features of wt nodules (Fig. 3), and that these were strongly repressed by N fertilization, together with their respective precursors choline, Pro, and nicotinic acid (Fig. 3c). Betaines are known as general osmoprotectants in prokaryotes and eukaryotes (Vriezen et al., 2007; Chen and Murata, 2011) and are essential for functional nodules (Goldmann et al., 1994). Rhizobia can take up betaines such as stachydrine and catabolize them by enzymes encoded by the *stc* operon (Goldmann et al., 1991; Phillips et al., 1998). Hence, betaines may support rhizobia either as nutrients or as compatible solutes against biotic stress resulting from the intracellular lifestyle and from manipulation by the host (Poole and Ledermann, 2022). Thus, betaines provided by the host may represent an essential substrate and their withdrawal by the host may contribute to sanctioning of the bacterial partner, in particular under HN conditions.

### A role for purine metabolism in sanctioning?

Purine metabolism was very responsive to sanctioning, in particular to HN treatment, where the levels of hypoxanthine and guanine were reduced whereas their catabolites allantoin, allantoic acid, and ureidoglycine were increased relative to the controls (**Figs. 3, 6**). This may reflect either a symptom of senescence and degradation, or, alternatively, an activation of ureide-based metabolism, as occurs in ureidic legumes (Tajima et al., 2004), in contrast to temperate legumes, such as *M. truncatula,* that transport their fixed N in the form of Asn (Snapp and Vance, 1986). Since ureides were detected exclusively in HN plants, this condition appears to entail unique metabolic changes opposite to the conditions resulting from lack of N-fixation (*nifH*, argon treatments).

### Central metabolism and sanctioning

A general finding of our study was the notion that changes in central metabolism were poorly matched by changes in the proteins related to these pathways (compare **Figs. 2-7** with **Tables S9-S14**). This contrasts with defense markers that were prominently induced both at the metabolomic and the proteomic levels (**Fig. 2,5; Table S1**). This poor correlation may be a sign of post-translational regulation; this is indeed the case for primary metabolism, where the activity of many enzymes is regulated by phosphorylation in prokaryotes and eukaryotes (Chandel, 2021; Gruber et al., 2021). Phosphoproteomic analysis revealed important regulators of sink-source interactions such as the plastidial triosephosphate/phosphate translocator, sucrose-cleaving enzymes such as invertase and sucrose synthase, and the glycolytic key enzyme fructose-bisphosphate aldolase, which were all among the hundred top hits in >13,000 peptides phosphorylated under sanctioning conditions (**Table S16**). Future work should address the functional relevance of the observed phosphorylation sites for the activity of these proteins in C metabolism and transport, and ultimately in sanctioning of bacterial cheaters.

### Changes in cellular dynamics associated with sanctioning

Secretion, membrane dynamics, and cytoskeletal reorganization are central for plant endosymbiosis with fungi and bacteria (Harrison, 2012; de Carvalho-Niebel et al., 2024). It is therefore not surprising that Exocyst components, ABC transporters, the membrane remodeling protein remorin, as well as the cytoskeletal proteins myosin XI and the 115 kD actin-bundling protein showed dynamic phosphorylation with significant increases under sanctioning conditions. Although the relevance of these phosphorylation events for symbiosis remains to be explored, it is interesting to note that the phosphorylated remorin belongs to a legume-specific class that is induced during symbiosis and therefore has been termed SymREM1 (Raffaele et al., 2007). It is the best-characterized member in a family of ten remorins in *M. truncatula* (Raffaele et al., 2007). Phosphorylations of *M. truncatula* SymREM1 observed during sanctioning coincided with the intrinsically disordered N-terminal part that is phosphorylated by the symbiosis receptor kinases NFR1 and SYMRK in *Lotus japonicus* levels of conservation (Marín and Ott, 2012), the phosphorylated residue Ser48 of *M. truncatula* is conserved in *L. japonicus* (**Fig. 10**), corresponding in this case to Ser44 which is phosphorylated *in vitro* by NFR1 and SYMRK (Toth et al., 2012). In *M. truncatula*, SymREM1 controls stability and turnover of the symbiosis receptor LYK3 at the plasma membrane, and is required for symbiotic signaling during infection (Lefebvre et al., 2010). It is therefore tempting to speculate that phosphorylation of *M. truncatula* SymREM1 during sanctioning could interfere with symbiotic signaling, thereby de-repressing defense mechanisms that are normally attenuated by symbiotic signaling, and which can become induced during senescence, or in the context of defective symbiotic communication (Berrabah et al., 2024).

**Fig. 10.**
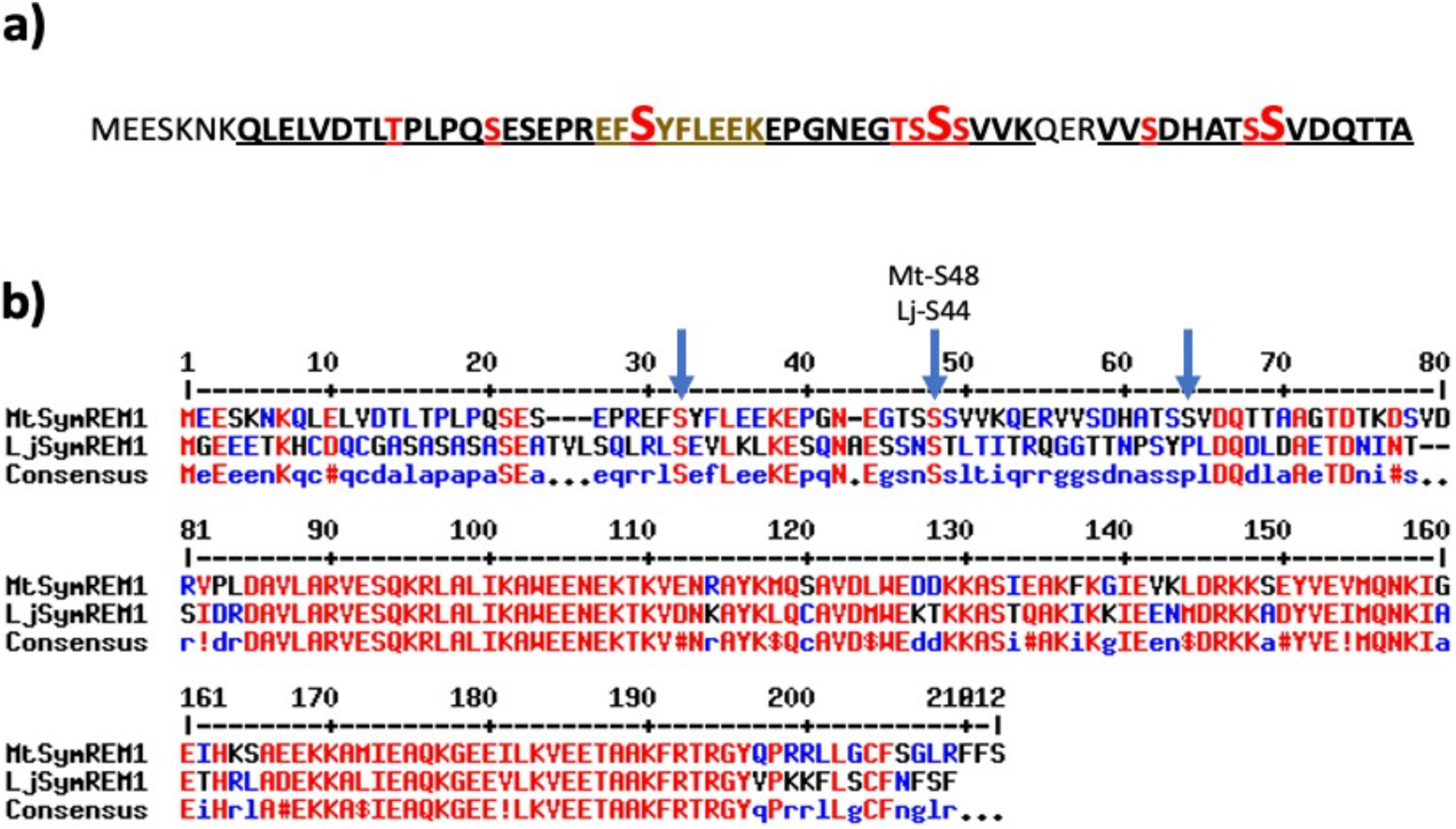
Phosphorylation sites of *M. truncatula* and *L. japonicus* SymREM1. **(a)** Phosphosites in the N-terminal intrinsically disordered region of *M. truncatula* SymREM1 are highlighted in red. Highly induced phosphorylation levels among the top 100 phosphopeptides (Table S16) are highlighted with increased font size. **(b)** Alignment of *M. truncatula* and *L. japonicus* SymREM1 amino acid sequence showing the N-terminal intrinsically disordered domain of *c.* 80 amino acids, which shows a very low level of sequence conservation, although two of the three phosphorylated Ser residues are conserved (arrows), in particular Ser44 which is phosphorylated by NFR1 and SYMRK in *L. japonicus* (Toth et al., 2012).

Finally, among the top 100 highly phosphorylated phosphopeptides were also six that represented two CCCH zinc-finger proteins, one represented by 5 phosphopeptides (**Table S16**). CCCH proteins occur in large families in various eukaryotes, and in plants, they are implicated in abiotic stress resistance and hormonal regulation (Han et al., 2021; Wang et al., 2022). Functional analysis using mutants of the genes encoding these yet unexplored proteins will reveal their roles in nodule symbiosis and sanctioning.

## Conclusion

Our three-level analysis of metabolome, proteome, and phosphoproteome revealed large scale transitions in host metabolism associated with sanctioning. Important individual symptoms of sanctioning, such as downregulation of NCRs and Lbs, and phosphorylation of RIN4, are each known to be sufficient to interfere with symbiosis. Thus, sanctioning is likely to represent a wide syndrome of anti-symbiotic effects, as has been evidenced by the inhibition of arbuscular mycorrhiza at high phosphate levels (Breuillin et al., 2010). Future research should address the contribution of phenomena that have not been functionally addressed, including changes in betaine and purine metabolism.

## Materials and methods

### Cultivation of rhizobia

*Sinorhizobium meliloti* strain 2011 was cultured in TY medium (5 g tryptone, 3 g yeast extract and 0.4 g CaCl_2_ in 1 liter distilled water) for recovery, in the presence of streptomycin (50 μg/ml), and incubated for 48 h. For *S. meliloti 2011 nifH*:Tn5, additional neomycin (50 μg/ml) was provided in TY medium, and incubated for 48 h.

### Plant growth, inoculation, and sanctioning treatments

Seeds of *M. truncatula* A17 were scarified by concentrated sulfuric acid for 10 min. After washing with sterile water three times, seeds were sterilized with concentrated Clorox (Reactolab SA) for 2 min, adding an equal volume of water for 1 min and rinsed with sterile water five times. Seeds were put in the shaker for 4 h. After soaking up water, seeds were put in 0.8% plant agar plates for 3 days. Five seedlings were transferred to one sterilized box unit (two sterilized cylindrical plastic boxes were linked by a cotton wick with perlite in the upper jar and B&D nutrient solution with 0.5 mM KNO_3_, Appendix 9) in the lower jar (Broughton and Dilworth, 1971). After two weeks, the plants were inoculated with strains *Sinorhizobium meliloti* 2011 or *S. meliloti* 2011 *nifH*:Tn5. Rhizobia were centrifuged at 4500 rpm for 15 min under 4℃ and re-suspended in 10 mM MgSO_4_. The OD600 of bacterial suspension was adjusted with 10 mM MgSO_4_ to reach about 0.2. Each plant was inoculated with 2 ml of bacterial suspension.

### RNA extraction and quantitative real-time RT-PCR for lignin biosynthetic genes

After the 21-day-treatment, nodules from *M. truncatula* were collected to extract total RNA. Six replicates were used for each treatment, and the nodules were ground into a fine powder using liquid nitrogen. Total RNA was extracted from the nodule powder following the protocol provided by the Spectrum™ Plant Total RNA-Kit (Sigma, https://www.sigmaaldrich.com). The total RNA was then converted to cDNA by reverse transcription with the SensiFAST™ cDNA Synthesis Kit (Bioline, https://www.bioline.com). For each quantitative real-time RT-PCR reaction, *c.* 10 ng of cDNA was mixed with 7.5 µl of FastStart Universal SYBR Green Master (Rox) (Roche, https://lifescience.roche.com), 0.5 µl of both forward and reverse primers (primer concentration 10 µM) (Appendix 7), and sterilized milliQ water to a total volume of 15 µl. qRT-PCR analysis was performed as follows: Start phase: 95℃ for 10 minutes; melting: 95℃ for 20 seconds; Annealing: 60℃ for 20 seconds; Extension:72℃ for 20 seconds. The constitutively genes *Mtc27* (TC132510) and *A39* were adopted as reference genes. Relative expression values were calculated and analyzed by ΔCt and one-way anova (Pfaffl, 2001; Pierre et al., 2014).

### Metabolite profiling on *Medicago truncatula* nodules

For metabolite profiling, inoculated plants were first grown under control conditions for two weeks, followed by treatments for 1 week under three sanctioning conditions, *nifH*, argon, and HN. Following the inoculation, plants were grown in open air for 14 days and then transferred to sealed boxes. Six jars inoculated with the *wt* rhizobial strain were treated with an argon mixture (80% argon, 20% O_2_, 400 ppm CO2), while the remaining plants were treated with air. After seven days of treatment, 100 mg nodules were harvested and pooled for each replicate pot. These nodules were subsequently ground into fine powder under liquid nitrogen. To this powder, 1 ml of 80% acetonitrile was added, and the nodule powder solution was sonicated twice, with each round lasting for 1 min. After letting the solution sit for 5 minutes on ice, the samples were centrifuged at 12,000 rpm for 3 min. The resulting supernatant (max 800 µl) was then collected for liquid chromatography and mass spectrometry analysis. Metabolites in the samples were separated by hydrophilic interaction liquid chromatography (HILIC) and analyzed after both positive and negative ionisation (see Supplemental Materials and Methos for further details). The data were analyzed using the MzMine2 program. Pure standard solutions of the compound of interest were injected for reference to verify predicted molecular identity by analysis of retention times and mass spectra was performed. Statistical analysis involved one-way analysis of variance (ANOVA) followed by Tukey’s test (p<0.05).

### Proteome sample preparation (label-free quantification)

Protein extraction was done according to Marx *et al*. (2016). Briefly, 100 mg of plant material was ground in liquid nitrogen to a fine powder. One ml of extraction buffer, 290 mM sucrose, 250 mM Tris-Cl, pH 7.6, protease inhibitor cocktail was added, well mixed, and sonicated. After removing of cell debris by centrifugation, one volume of chloroform was added and samples were mixed well. After adding 3 volumes of water and mixing, sample was centrifuged 5 min at 14,000g at 4°C. The upper phase was removed and 3 volumes of methanol was added to the lower phase and interphase and mixed well. After centrifugation, the pellet was washed 3 times with 80% acetone and dried at RT. Pellets were resuspended in 8 M urea in 50 mM Tris-Cl, pH 8, reduced with DTT (final concentration 1 mM, 30 min at RT) and alkylated with IAA (final concentration 5.5 mM, 30 min at RT in the dark). Afterwards, the same amount of proteins per sample was digested with LysC (ratio 50:1, for 3 h, at RT, final urea concentration 4 M) and trypsin (ratio 50:1, overnight, at RT, final urea concentration <1 M). Next day, the samples were acidified using 50% TFA (final concentration app. 0.5%, pH<2) and centrifuged at 4000 rpm for 10 min to remove precipitations. Peptides were purified by SPE using HR-X columns in combination with C18 cartridges (Macherey-Nagel; https://www.mn-net.com): wash buffer, 0.1% formic acid in deionized water; elution buffer, 80% acetonitrile, and 0.1% formic acid in deionized water. Elutes were frozen in liquid nitrogen and lyophilized overnight. Purified peptides were fractionated by HpH reversed phase chromatography (Batth *et al*., 2014). Briefly, dry peptide powder was suspended in 400 μl 5% ammonium hydroxide and fractionated using a Waters XBridge BEH130 C18 column (3.5 μm 4.6 × 250 mm) on a Ultimate 3000 HPLC (Thermo Scientific). The flow rate of the mobile phase was 1 ml/min. HpH buffer A contained 10 mM ammonium formate in deionized water and buffer B contained 10 mM ammonium formate and 90% acetonitrile deionized water. Both buffers were adjusted to pH 10 with ammonium hydroxide. Peptide fractions were acidified, frozen in liquid nitrogen, and lyophilized overnight.

LC-MS/MS measurements were performed on an QExactive HFX mass spectrometer coupled to an EasyLC 1000 nanoflow-HPLC (all Thermo Scientific). Peptides were separated on a fused silica HPLC-column tip (I.D. 75 μm, New Objective, self-packed with ReproSil-Pur 120 C18-AQ, 1.9 μm (Dr. Maisch) to a length of 20 cm) using a gradient of A (0.1% formic acid in water) and B (0.1% formic acid in 80% acetonitrile in water): samples were loaded with 0% B with a flow rate of 600 nl/min; peptides were separated by 5%–30% B within 85 min with a flow rate of 250 nl/min. Spray voltage was set to 2.3 kV and the ion-transfer tube temperature to 250°C; no sheath and auxiliary gas were used. Mass spectrometers were operated in the data-dependent mode; after each MS scan (*m/z* = 370 – 1750; resolution: 120,000) a maximum of twelve MS/MS scans were performed using an isolation window of 1.6, a normalized collision energy of 28%, a target AGC of 1e5 and a resolution of 30,000. MS raw files were analyzed using MaxQuant software using a Uniprot full-length *M. truncatula* database and a *S. meliloti* database. Carbamidomethylcysteine was set as fixed modification and protein amino-terminal acetylation and oxidation of methionine were set as variable modifications. The MS/MS tolerance was set to 20 ppm and three missed cleavages were allowed using trypsin/P as enzyme specificity. Peptide and protein FDR based on a forward-reverse database were set to 0.01, minimum peptide length was set to 7, and minimum number of peptides for identification of proteins was set to one, which must be unique. The “match-between-run” option was used with a time window of 0.7 min. MaxQuant results were analyzed using Perseus.

### Proteome sample preparation (tandem mass tag-based quantification)

After extraction of metabolites, samples were lyophilized before resuspending pellet in 6 M guanidine hydrochloride (GdnHCl) in 100 mM HEPES, pH 8. After mixing, samples were sonicated and cell debris were removed by centrifugation. Proteins were reduced with TCEP (final concentration 10 mM, 1 h at 55°C) and alkylated with IAA (final concentration 22 mM, 30 min at RT). After acetone precipitation, proteins were resuspended in small volume of 2.5 M GdnHCl in 100 mM HEPES, pH 8 and 150 μg protein of each sample were digested overnight with LysC and trypsin (ratio 50:1, final volume of 100 μl, final GdnHCl concentration of 0.75 M). TMTpro labeling was performed according to the manufacturer’s recommendations. After combining same amount of each TMT channel, peptide mixture was fractionated by HpH reverse phase chromatography (see above) resulting in eight fractions. Phosphopeptide enrichment was done according to Post *et al*. (2017) using Fe(III) cartridges on Bravo liquide handling system (Agilent).

LC-MS/MS measurements were performed on an Exploris 480 mass spectrometer coupled to an EasyLC 1200 nanoflow-HPLC (all Thermo Scientific). Peptides were seperated on a fused silica HPLC-column tip (I.D. 75 μm, New Objective, self-packed with ReproSil-Pur 120 C18-AQ, 1.9 μm (Dr. Maisch) to a length of 20 cm) using a gradient of A (0.1% formic acid in water) and B (0.1% formic acid in 80% acetonitrile in water): samples were loaded with 0% B with a flow rate of 600 nl/min; peptides were separated by 7%–38% B within 122 min with a flow rate of 250 nl/min. Spray voltage was set to 2.3 kV and the ion-transfer tube temperature to 250°C; no sheath and auxiliary gas were used. Mass spectrometers were operated in the data-dependent mode; after each MS scan ( *m/z* = 370– 1750; resolution: 120,000) a maximum of twenty MS/MS scans were performed using an isolation window of 0.7, a normalized collision energy of 32%, a target AGC of 50% and a resolution of 45,000. MS raw files were analyzed using ProteomeDiscoverer (version 2.5, Thermo Scientific) using a Uniprot full-length *Medicago truncatula* database and a *Sinorhizobium meliloti* database. Carbamidomethylcysteine was set as fixed modification and protein amino-terminal acetylation and oxidation of methionine were set as variable modifications. The MS/MS tolerance was set to 0.6 Da and three missed cleavages were allowed using trypsin/P as enzyme specificity. TMTpro correction factors were used. Exported normalized abundances for each channel were further analyzed using Perseus software (Tyanova *et al*., 2016).

## Acknowledgements

We thank Peter Mergaert for providing the bacterial strains and Bettina Hause for *M. truncatula* seeds.

## Author contributions

MC, AR, and NJ performed experiments; PMA, ED and MS performed metabolomic and proteomic analysis; IY and MB carried out protein 3D modeling and phosphoproteome analysis; DR, MC, and AR wrote the manuscript; DR conceived and supervised the project.

