## Supplemental Figures and Tables 1,2,3,7,8 for "Sanctioning of bacterial cheaters by the host plant in nitrogen-fixing symbiosis between *Medicago truncatula* and *Sinorhizobium meliloti*"

### SUPPLEMENTARY FIGURES

**Fig. S1. Uric acid / Purin degradation pathway**

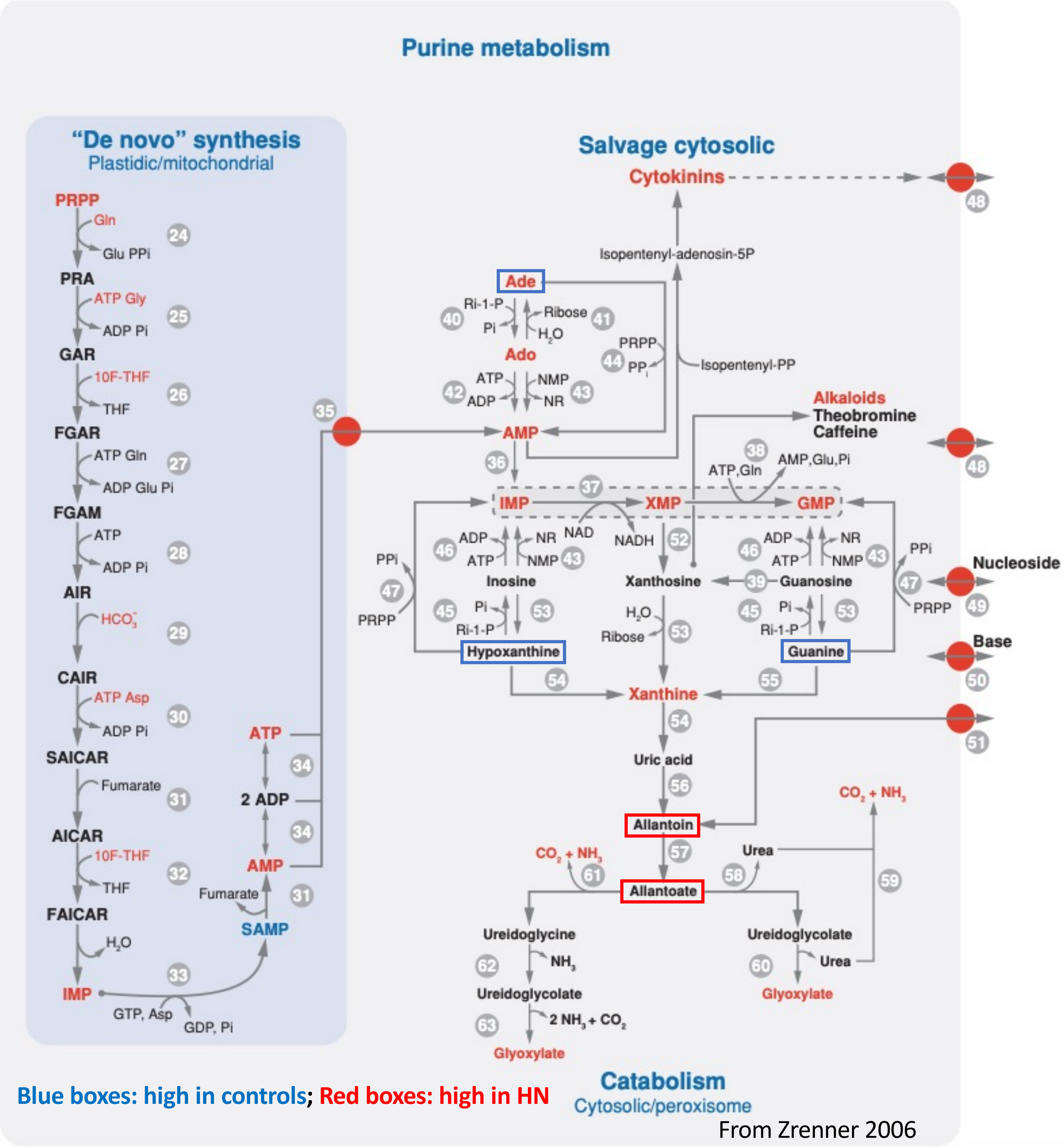

Blue boxes: high in controls; Red boxes: high in HN

From Zrenner 2006

**Fig. S2. Proteomic experiment 1: Global clustering**

Free-label significant ID both in nifH and argon samples

Total protein ID number 558

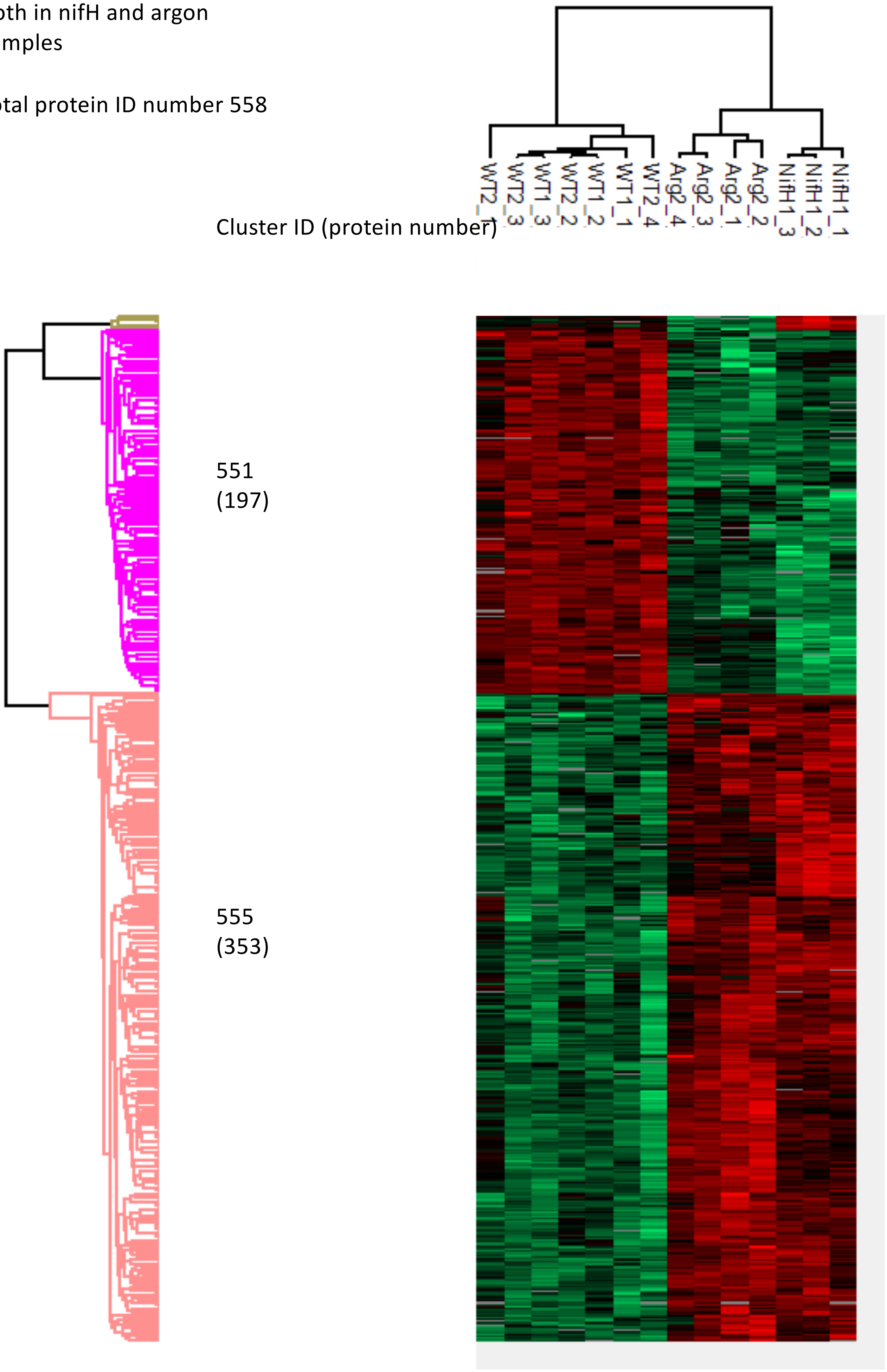

**Fig. S3. Proteomic experiment 2: Global clustering**

TMT significant ID all in nifH argon and HN samples

Total protein ID number 1097

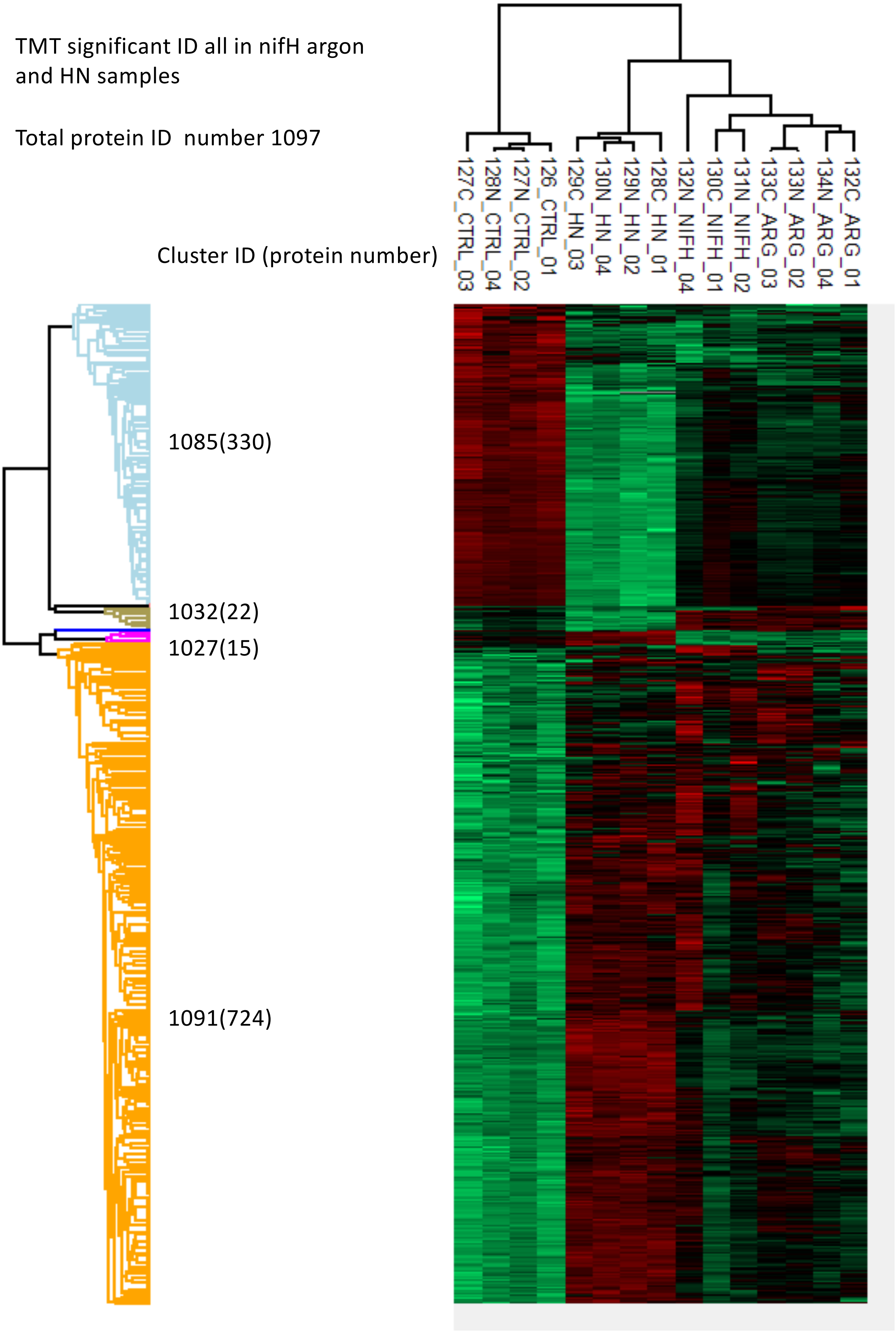

**Table S1**

### Up-regulated GO enrichment

**a)**

| | nifH-wt | | | Argon-control | | | |
|---|---|---|---|---|---|---|---|
| | Count | Expected | q-value | Count | Expected | q-value | Aspect |
| single-organism catabolic process | 38 | 15.91 | 0.00339 | 58 | 17.34 | 8.88E-13 | P |
| single-organism metabolic process | 222 | 168.69 | 0.00436 | 307 | 183.83 | 2.03E-20 | P |
| immune response * | 21 | 6.73 | 0.00646 | 24 | 7.33 | 5.3E-05 | P |
| innate immune response * | 20 | 6.55 | 0.00692 | 23 | 7.13 | 0.000102 | P |
| defense response to fungus, incompatible interaction * | 8 | 1.19 | 0.00872 | 10 | 1.3 | 5.3E-05 | P |
| response to inorganic substance | 40 | 19.67 | 0.00872 | 52 | 21.44 | 9.96E-07 | P |
| immune system process * | 21 | 7.52 | 0.00894 | 25 | 8.2 | 0.000101 | P |
| response to external biotic stimulus | 39 | 19.2 | 0.00894 | 53 | 20.92 | 2.14E-07 | P |
| response to other organism | 39 | 19.2 | 0.00894 | 53 | 20.92 | 2.14E-07 | P |
| defense response to other organism * | 32 | 14.54 | 0.00904 | 39 | 15.84 | 3.95E-05 | P |
| defense response, incompatible interaction * | 14 | 3.83 | 0.00904 | 18 | 4.18 | 2.98E-05 | P |
| small molecule catabolic process | 14 | 3.94 | 0.011 | 23 | 4.3 | 2.74E-08 | P |
| response to biotic stimulus | 41 | 21.05 | 0.011 | 58 | 22.93 | 5.81E-08 | P |
| lipid catabolic process | 13 | 3.47 | 0.011 | 16 | 3.78 | 0.000127 | P |
| single-organism process | 392 | 341.14 | 0.0114 | 408 | 371.75 | 0.1435 | P |
| response to alcohol | 28 | 13.13 | 0.0271 | 26 | 14.3 | 0.1096 | P |
| response to external stimulus | 45 | 25.42 | 0.0271 | 64 | 27.7 | 1.89E-07 | P |
| benzene-containing compound metabolic process | 6 | 0.87 | 0.0271 | 6 | 0.95 | 0.01816 | P |
| response to fungus * | 18 | 7.2 | 0.0457 | 26 | 7.84 | 1.61E-05 | P |
| organic acid catabolic process | 10 | 2.75 | 0.0495 | 18 | 2.99 | 2.18E-07 | P |
| carboxylic acid catabolic process | 10 | 2.75 | 0.0495 | 18 | 2.99 | 2.18E-07 | P |
| cinnamic acid biosynthetic process | 3 | 0.18 | 0.0495 | 4 | 0.2 | 0.001237 | P |
| cinnamic acid metabolic process | 3 | 0.18 | 0.0495 | 4 | 0.2 | 0.001237 | P |
| response to abscisic acid | 24 | 11.32 | 0.0495 | 24 | 12.33 | 0.07582 | P |

Ordered according to p-values in nifH-wt

| | | | | |
|---|---|---|---|---|
| 0 | 0.01 | 0.03 | 0.05 | 1 |

**Table S2**

lab meeting20211221

### Down-regulated GO enrichment

**b)**

| | nifH-wt | | | Argon-control | | | |
|---|---|---|---|---|---|---|---|
| | Count | Expected | q-value | Count | Expected | q-value | Aspect |
| nodulation * | 39 | 7.53 | 1.07E-13 | 37 | 8.27 | 8.072E-11 | P |
| interspecies interaction between organisms | 40 | 8.1 | 1.07E-13 | 37 | 8.9 | 1.84E-10 | P |
| symbiosis, encompassing mutualism through parasitism * | 39 | 8.06 | 3.71E-13 | 37 | 8.86 | 1.84E-10 | P |
| nodule morphogenesis * | 37 | 7.33 | 3.97E-13 | 36 | 8.06 | 8.072E-11 | P |
| development involved in symbiotic interaction * | 37 | 7.35 | 3.97E-13 | 36 | 8.07 | 8.072E-11 | P |
| post-embryonic morphogenesis | 38 | 8.83 | 2.02E-11 | 37 | 9.7 | 1.937E-09 | P |
| anatomical structure morphogenesis | 45 | 14.22 | 2.91E-09 | 40 | 15.63 | 0.00002698 | P |
| multi-organism process | 46 | 18.19 | 2.36E-06 | 45 | 19.99 | 0.0001271 | P |
| single-organism process | 158 | 114.54 | 2.64E-06 | 156 | 125.89 | 0.02724 | P |
| post-embryonic development | 46 | 18.65 | 4.12E-06 | 45 | 20.5 | 0.0002314 | P |
| anatomical structure development | 56 | 25.68 | 5.28E-06 | 51 | 28.22 | 0.007078 | P |
| single-multicellular organism process | 55 | 26.13 | 2.26E-05 | 53 | 28.72 | 0.00339 | P |
| single-organism developmental process | 57 | 27.73 | 2.53E-05 | 54 | 30.48 | 0.007078 | P |
| multicellular organismal development | 54 | 25.69 | 2.80E-05 | 52 | 28.24 | 0.004095 | P |
| developmental process | 57 | 28.15 | 3.55E-05 | 54 | 30.95 | 0.009686 | P |
| multicellular organismal process | 55 | 28.37 | 2.54E-04 | 54 | 31.19 | 0.0113 | P |
| glutamine family amino acid biosynthetic process | 6 | 0.39 | 5.98E-04 | 3 | 0.43 | 1 | P |
| nitrogen fixation * | 2 | 0.02 | 3.16E-02 | 2 | 0.03 | 0.05128 | P |
| gas transport | 2 | 0.04 | 6.87E-02 | 2 | 0.04 | 0.1351 | P |
| oxygen transport * | 2 | 0.04 | 6.87E-02 | 2 | 0.04 | 0.1351 | P |
| microtubule-based process | 8 | 1.88 | 9.38E-02 | 8 | 2.07 | 0.2785 | P |
| asparagine metabolic process | 2 | 0.06 | 1.87E-01 | 2 | 0.07 | 0.3808 | P |
| asparagine biosynthetic process | 2 | 0.06 | 1.87E-01 | 2 | 0.07 | 0.3808 | P |
| glutamine biosynthetic process | 2 | 0.13 | 6.50E-01 | 2 | 0.15 | 0.00901 | P |

Ordered according to p-values in nifH-wt

| | | | | |
|---|---|---|---|---|
| 0 | 0.01 | 0.03 | 0.05 | 1 |

**Fig. S4. induced defense markers in nifH and argon nodules**

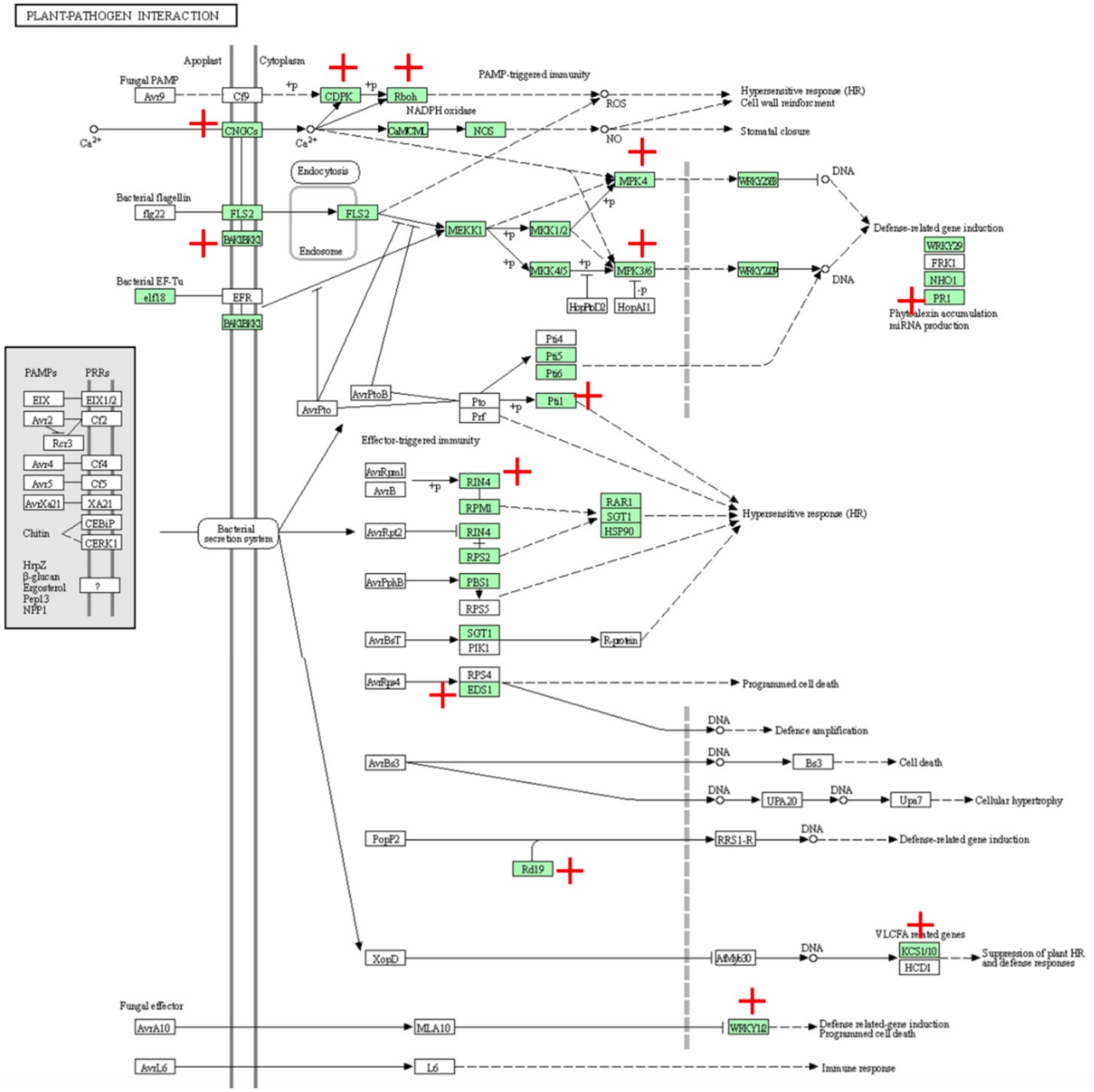

### Fig. S5. GO terms overrepresented in argon & nifH & HN vs. controls (TMT-labelled)

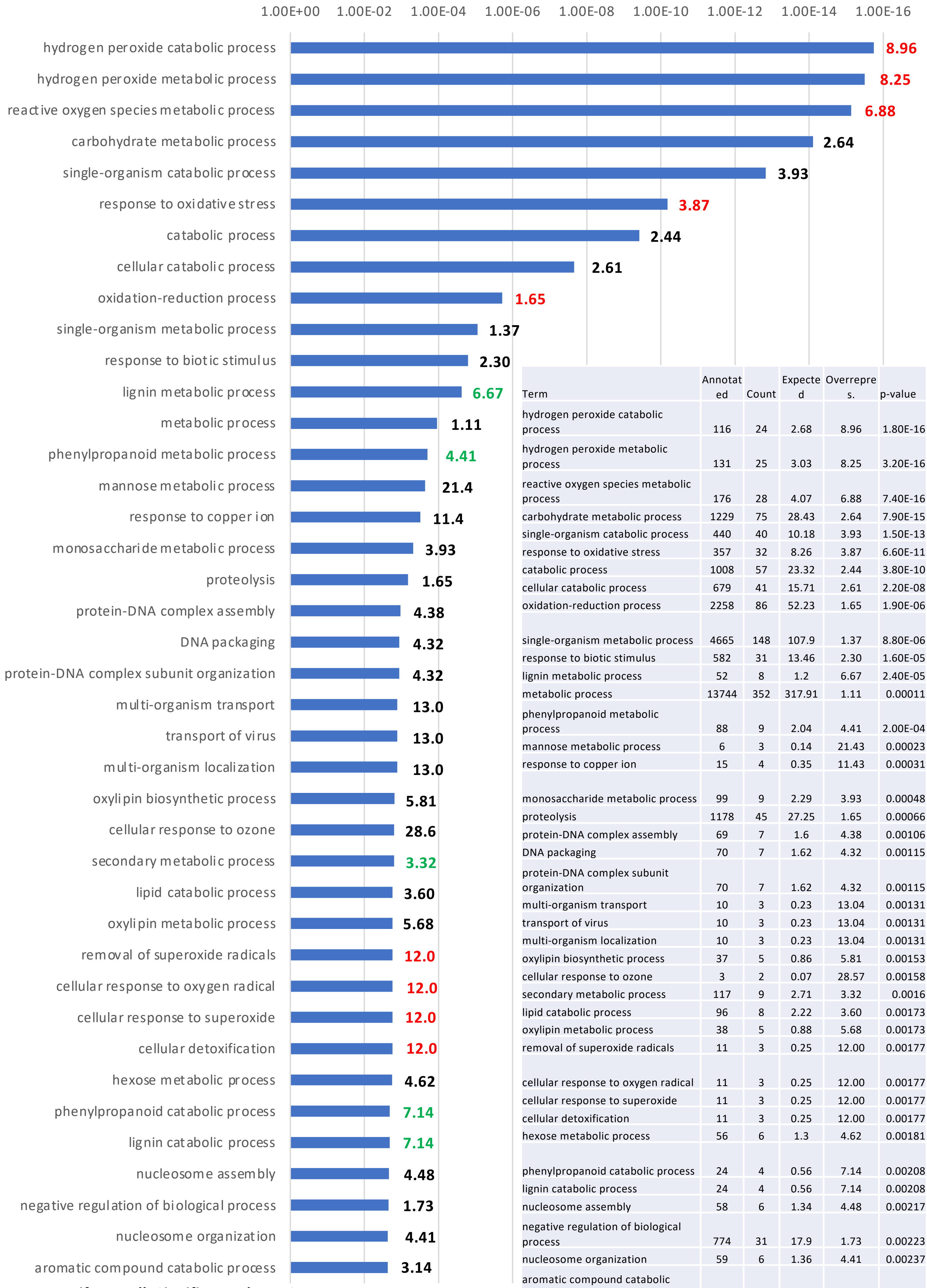

| Term | Annotated | Count | Expected | Overrepres. | p-value |
|---|---|---|---|---|---|
| hydrogen peroxide catabolic process | 116 | 24 | 2.68 | 8.96 | 1.80E-16 |
| hydrogen peroxide metabolic process | 131 | 25 | 3.03 | 8.25 | 3.20E-16 |
| reactive oxygen species metabolic process | 176 | 28 | 4.07 | 6.88 | 7.40E-16 |
| carbohydrate metabolic process | 1229 | 75 | 28.43 | 2.64 | 7.90E-15 |
| single-organism catabolic process | 440 | 40 | 10.18 | 3.93 | 1.50E-13 |
| response to oxidative stress | 357 | 32 | 8.26 | 3.87 | 6.60E-11 |
| catabolic process | 1008 | 57 | 23.32 | 2.44 | 3.80E-10 |
| cellular catabolic process | 679 | 41 | 15.71 | 2.61 | 2.20E-08 |
| oxidation-reduction process | 2258 | 86 | 52.23 | 1.65 | 1.90E-06 |
| single-organism metabolic process | 4665 | 148 | 107.9 | 1.37 | 8.80E-06 |
| response to biotic stimulus | 582 | 31 | 13.46 | 2.30 | 1.60E-05 |
| lignin metabolic process | 52 | 8 | 1.2 | 6.67 | 2.40E-05 |
| metabolic process | 13744 | 352 | 317.91 | 1.11 | 0.00011 |
| phenylpropanoid metabolic process | 88 | 9 | 2.04 | 4.41 | 2.00E-04 |
| mannose metabolic process | 6 | 3 | 0.14 | 21.43 | 0.00023 |
| response to copper ion | 15 | 4 | 0.35 | 11.43 | 0.00031 |
| monosaccharide metabolic process | 99 | 9 | 2.29 | 3.93 | 0.00048 |
| proteolysis | 1178 | 45 | 27.25 | 1.65 | 0.00066 |
| protein-DNA complex assembly | 69 | 7 | 1.6 | 4.38 | 0.00106 |
| DNA packaging | 70 | 7 | 1.62 | 4.32 | 0.00115 |
| protein-DNA complex subunit organization | 70 | 7 | 1.62 | 4.32 | 0.00115 |
| multi-organism transport | 10 | 3 | 0.23 | 13.04 | 0.00131 |
| transport of virus | 10 | 3 | 0.23 | 13.04 | 0.00131 |
| multi-organism localization | 10 | 3 | 0.23 | 13.04 | 0.00131 |
| oxylipin biosynthetic process | 37 | 5 | 0.86 | 5.81 | 0.00153 |
| cellular response to ozone | 3 | 2 | 0.07 | 28.57 | 0.00158 |
| secondary metabolic process | 117 | 9 | 2.71 | 3.32 | 0.0016 |
| lipid catabolic process | 96 | 8 | 2.22 | 3.60 | 0.00173 |
| oxylipin metabolic process | 38 | 5 | 0.88 | 5.68 | 0.00173 |
| removal of superoxide radicals | 11 | 3 | 0.25 | 12.00 | 0.00177 |
| cellular response to oxygen radical | 11 | 3 | 0.25 | 12.00 | 0.00177 |
| cellular response to superoxide | 11 | 3 | 0.25 | 12.00 | 0.00177 |
| cellular detoxification | 11 | 3 | 0.25 | 12.00 | 0.00177 |
| hexose metabolic process | 56 | 6 | 1.3 | 4.62 | 0.00181 |
| phenylpropanoid catabolic process | 24 | 4 | 0.56 | 7.14 | 0.00208 |
| lignin catabolic process | 24 | 4 | 0.56 | 7.14 | 0.00208 |
| nucleosome assembly | 58 | 6 | 1.34 | 4.48 | 0.00217 |
| negative regulation of biological process | 774 | 31 | 17.9 | 1.73 | 0.00223 |
| nucleosome organization | 59 | 6 | 1.36 | 4.41 | 0.00237 |
| aromatic compound catabolic process | 124 | 9 | 2.87 | 3.14 | 0.00238 |

**TMT_Arg NifH HN all_Significant_cluster2**

**Table S3. Levels of proteins identified with search terms related to defense**

| | | Free-label | | TMT-label | | | |
|---|---|---|---|---|---|---|---|
| Family | Uniprot | nifH/wt | argon/control | Arg/Ctrl | HN/Ctrl | NifH/Ctrl | description |
| CDPK | | | | | | | Calcium-dependent protein kinase |
| | G7I8E2 | 1.46 | 1.14 | 0.89 | 1.11 | 1.33 | Calcium-dependent kinase |
| | A0A072VTY6 | 1.84 | 1.29 | 1.07 | 1.07 | 1.06 | Calmodulin-domain kinase CDPK protein |
| | A0A072UWP5 | 1.47 | 1.65 | 1.17 | 1.40 | 1.12 | Calmodulin-domain kinase CDPK protein |
| | A0A072V1D7 | 1.50 | 1.30 | 1.17 | 1.23 | 1.08 | Calmodulin-domain kinase CDPK protein |
| | G7K8K5 | 1.21 | 1.36 | 1.10 | 1.33 | 1.13 | Calcium-dependent kinase |
| | A0A072UFA6 | 1.21 | 1.36 | | | | Calcium-dependent kinase |
| | A0A072U1R9 | 1.46 | 1.47 | 1.20 | 1.32 | 1.14 | Calmodulin-domain kinase CDPK protein |
| CNGCs | | | | | | | cyclic nucleotide gated channel |
| | A0A072VMJ3 | 1.40 | 1.01 | | | | Cyclic nucleotide-gated channel involved in the establishment of both rhizobial and m |
| | G7J6T3 | 1.56 | 1.02 | | | | Cyclic nucleotide-gated ion channel-like protein |
| | G7JND3 | 1.40 | 1.01 | 1.09 | 0.93 | 1.09 | Cyclic nucleotide-gated channel involved in the establishment of both rhizobial and m |
| | G7JGP4 | 1.40 | 1.01 | | | | Cyclic nucleotide-gated ion channel-like protein |
| | A0A072U5G5 | nd | 1.88 | | | | Cyclic nucleotide-gated ion channel-like protein |
| Rboh | | | | | | | respiratory burst oxidase |
| | G7I8P3 | 0.36 | 0.57 | 0.89 | 0.71 | 0.69 | Putative NAD(P)H oxidase (H(2)O(2)-forming) |
| | G7J2Q7 | 1.66 | 1.43 | | | | Putative NAD(P)H oxidase (H(2)O(2)-forming) |
| | G7J2Q9 | nd | 5.42 | | | | Putative NAD(P)H oxidase (H(2)O(2)-forming) |
| | Q5ENY3 | 1.44 | 1.53 | 1.24 | 1.28 | 1.10 | Calcium-binding EF-hand; Ferric reductase-like transmembrane component |
| | G7LGI5 | 1.31 | 1.38 | | | | Respiratory burst oxidase-like protein D |
| | A0A072U5M5 | 0.44 | 0.39 | | | | Putative NAD(P)H oxidase (H(2)O(2)-forming) |
| BAK-IRK | | | | | | | brassinosteroid insensitive 1-associated receptor kinase 1 |
| | E2IXG1 | 1.46 | 1.31 | 1.10 | 1.35 | 1.17 | Non-specific serine/threonine protein kinase |
| | G7ILB9 | 1.46 | 1.31 | | | | Non-specific serine/threonine protein kinase |
| MPK4 | | | | 1.30 | 1.66 | 1.04 | Mitogen-activated protein kinase 4 |
| | G7L6L0 | 1.06 | 1.44 | 1.13 | 1.42 | 1.05 | Mitogen-activated protein kinase |
| | A0A072TMU3 | 1.16 | 1.31 | | | | Mitogen-activated protein kinase |
| MPK3/6 | | | | | | | Mitogen-activated protein kinase 3 |
| | G7JNP9 | 1.33 | 1.13 | 1.12 | 1.35 | 1.05 | Mitogen-activated protein kinase |
| | G7JRI4 | 1.46 | 1.49 | 1.08 | 1.19 | 0.98 | Mitogen-activated protein kinase |
| PR1 | | | | | | | pathogenesis-related protein 1 |
| | G7IFD0 | 0.93 | 3.28 | 1.62 | 1.83 | 1.10 | CAP, cysteine-rich secretory protein, antigen 5 |
| | G7ILE4 | 0.93 | 3.28 | | | | CAP, cysteine-rich secretory protein, antigen 5 |
| | I3SBC6 | 1.69 | 4.17 | 1.49 | 1.67 | 1.06 | CAP, cysteine-rich secretory protein, antigen 5 |
| Pti1 | | | | | | | pto-interacting protein 1 |
| | G7IBS1 | 1.27 | 1.09 | 1.06 | 1.13 | 1.04 | Receptor-like kinase |
| | G7IW36 | 1.87 | 1.84 | | | | NAD(P)H dehydrogenase (quinone)/PTI1-like tyrosine-kinase |
| | G7JQ35 | 1.69 | 1.80 | 1.20 | 1.39 | 1.26 | Pti1-like kinase |
| RIN4 | | | | | | | RPM1-interacting protein 4 |
| | B7FGR2 | 1.49 | 1.11 | | | | Putative RIN4, pathogenic type III effector avirulence factor Avr cleavage |
| | A0A072TXB9 | 1.97 | 1.63 | 1.39 | 1.48 | 1.28 | Putative RIN4, pathogenic type III effector avirulence factor Avr cleavage |
| RPM1 | | | | | | | disease resistance protein RPM1 |
| | G7I6E8 | 1.34 | 1.40 | 1.05 | 1.16 | 1.10 | NB-ARC domain disease resistance protein |
| | A0A072V485 | 1.42 | 1.56 | | | | NBS-LRR type disease resistance protein |
| | A0A072UT37 | 1.42 | 1.56 | 1.26 | 1.63 | 1.26 | Disease resistance protein (CC-NBS-LRR class) family protein |
| | A0A072UTM5 | nd | 1.48 | 0.84 | 1.21 | 1.22 | NBS-LRR type disease resistance protein |
| | G7J231 | 1.37 | 1.73 | | | | LRR and NB-ARC domain disease resistance protein |
| | G7J232 | 1.72 | 2.35 | | | | LRR and NB-ARC domain disease resistance protein |
| | G7J234 | 1.37 | 1.73 | | | | LRR and NB-ARC domain disease resistance protein |
| | A0A072V757 | 1.34 | 1.40 | 1.17 | 1.14 | 1.22 | NB-ARC domain disease resistance protein |
| | A0A072UY20 | nd | nd | | | | NB-ARC domain disease resistance protein |
| | A0A072UVY0 | 1.34 | 1.40 | | | | NB-ARC domain disease resistance protein |
| | G7K9I6 | | | | | | Disease resistance protein (CC-NBS-LRR class) family protein |
| | G7K9J0 | | | | | | Disease resistance protein (CC-NBS-LRR class) family protein |
| EDS1 | | | | | | | enhanced disease susceptibility 1 protein |
| | A0A072V056 | 1.11 | 1.67 | 1.34 | 1.82 | 1.19 | Enhanced disease susceptibility protein |
| Rd19 | | | | | | | cathepsin F |
| | G7I5V9 | 1.80 | 1.10 | 1.11 | 1.47 | 1.12 | Papain family cysteine protease |
| | G7JIL2 | 2.34 | 1.56 | 1.24 | 1.72 | 1.32 | Papain family cysteine protease |
| | G7JL32 | nd | nd | | | | Papain family cysteine protease |
| | A0A072UXB2 | 2.34 | 1.56 | | | | Papain family cysteine protease |
| KCS1/10 | | | | | | | 3-ketoacyl-CoA synthase |
| | G7ZZE4 | nd | nd | | | | 3-ketoacyl-CoA synthase |
| | G7J960 | nd | nd | | | | 3-ketoacyl-CoA synthase |
| | A0A072UVL7 | 1.38 | 1.34 | | | | 3-ketoacyl-CoA synthase |
| | G7JJR3 | 1.38 | 1.34 | | | | 3-ketoacyl-CoA synthase |
| | G7JHB4 | 1.38 | 1.34 | | | | 3-ketoacyl-CoA synthase |
| | G7K1J8 | nd | nd | | | | 3-ketoacyl-CoA synthase |
| | G7KJN5 | 1.38 | 1.34 | | | | 3-ketoacyl-CoA synthase |
| | Q2HTC9 | nd | nd | | | | 3-ketoacyl-CoA synthase |
| WRKY1/2 | | | | | | | WRKY transcription factor 2 |
| | G7KEQ1 | 1.55 | 1.09 | 1.32 | 1.12 | 1.24 | Putative transcription factor WRKY family |

#### Fig. S6. Lignin biosynthetic pathway

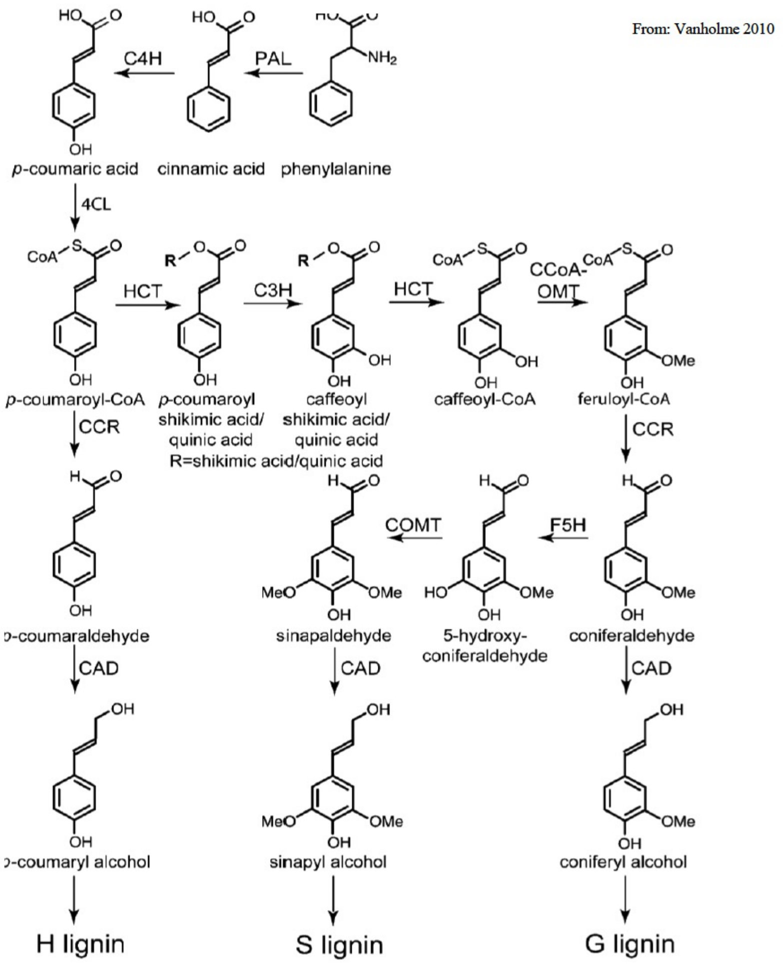

The main biosynthetic route toward the monolignols p-coumaryl, coniferyl, and sinapyl alcohol (Boerjan et al., 2003).
PAL, PHENYLALANINE AMMONIA-LYASE;
C4H, CINNAMATE 4-HYDROXYLASE;
4CL, 4-COUMARATE:CoA LIGASE;
C3H, p-COUMARATE 3-HYDROXYLASE;
HCT, p-HYDROXYCINNAMOYL-CoA:QUINATE/ SHIKIMATE p-HYDROXYCINNAMOYLTRANSFERASE;
CCoAOMT, CAFFEOYL-CoA O-METHYLTRANSFERASE;
CCR, CINNAMOYL-CoA REDUCTASE;
F5H, FERULATE 5-HYDROXYLASE;
COMT, CAFFEIC ACID O-METHYLTRANSFERASE;
CAD, CINNAMYL ALCOHOL DEHYDROGENASE.

#### Table S4. Expression of lignin biosynthetic genes under conditions of sanctioning

| Genes | Protein ID | Proteomics | | qRT-PCR | |
|---|---|---|---|---|---|
| | | nifH/wt | Argon/control | nifH/wt | Argon/control |
| PAL-1 | G7IBI3 | **1.75** | **2.65** | **30.71** | **14.72** |
| PAL-2 | G7I515 | **2.07** | **1.78** | 3.13 | 5.68 |
| C4H | Q2MJ09 | **2.14** | **1.69** | **14.92** | **12.04** |
| 4CL-1 | G7JVI7 | 1.16 | **2.50** | **6.22** | **4.79** |
| 4CL-2 | G7K6G3 | **1.73** | **2.45** | 1.35 | 1.91 |
| 4CL-3 | G7KG19 | nd | nd | **3.23** | **4.32** |
| HCT | G7L0F4 | **3.53** | **4.62** | 1.35 | 0.78 |
| C3H1 | A0A396H909 | nd | nd | **3.25** | **13.34** |
| CCoAOMT-1 | G7JK14 | 1.24 | **1.61** | **2.47** | **4.96** |
| CCoAOMT-2 | A0A072UQF3 | 0.76 | 0.90 | 0.45 | 1.65 |
| CCoAOMT-3 | A0A072UQ24 | **3.99** | **5.55** | **2.01** | **5.91** |
| CCR-1 | A0A072VDF2 | 1.09 | 1.20 | **2.77** | **8.06** |
| CCR-2 | G7JFN0 | 1.38 | **1.95** | 0.64 | 1.02 |
| F5H1 | G7JJV2 | nd | 0.94 | **1.64** | 1.15 |
| COMT-1 | A0A396J046 | nd | nd | **2.23** | 1.91 |
| COMT-2 | G7K511 | **1.48** | **2.33** | **0.58** | **2.32** |
| COMT-3 | A0A072VZR2 | **2.33** | **3.23** | 1.44 | **3.91** |
| CAD | A0A072VPZ8 | 1.21 | 1.63 | **2.17** | **3.75** |

#### Table S7. Relative leghemoglobin protein levels under conditions of sanctioning

| | | | continuos treatment | | 7 days treatment | | |
|---|---|---|---|---|---|---|---|
| | | | nifH /wt | argon/wt | nifH/wt | argon/air | HN/control |
| Leghemoglobin 1 | Medtr5g080440 | G7KGT0 | **0.57** | **0.64** | **0.64** | **0.52** | **0.19** |
| Leghemoglobin 2 | Medtr5g081030 | G7K1Z9 | 0.67 | **0.61** | **0.67** | **0.58** | **0.24** |
| Leghemoglobin 3 | Medtr5g080400 | G7KGS8 | **0.29** | **0.60** | **0.56** | **0.46** | **0.19** |
| Leghemoglobin 4 | Medtr1g049330 | A0A072VIZ0 | **0.58** | **0.63** | **0.78** | **0.52** | **0.23** |
| Leghemoglobin 5 | Medtr5g080900 | G7K1Y9 | nd | nd | nd | nd | nd |
| Leghemoglobin 6 | Medtr5g041610 | G7KGN2 | **0.44** | **0.43** | **0.60** | **0.53** | **0.25** |
| Leghemoglobin 7 | Medtr1g011540 | G7I6B5 | **0.44** | **0.52** | **0.56** | **0.51** | **0.26** |
| Leghemoglobin 8 | Medtr5g081000 | G7K1Z7 | **0.28** | **0.72** | **0.64** | **0.56** | **0.30** |
| Leghemoglobin 9 | Medtr5g066070 | G7KBK3 | **0.42** | **0.54** | **0.48** | **0.45** | **0.22** |
| Leghemoglobin 10 | Medtr1g090820 | A0A072VZ72 | 0.72 | 0.72 | **0.62** | **0.59** | **0.12** |
| Leghemoglobin 11 | Medtr1g090810 | B3SGL5 | **0.34** | **0.37** | **0.52** | **0.45** | **0.25** |
| Leghemoglobin 12 | Medtr7g110180 | G7KTR8 | nd | nd | nd | nd | nd |

#### Table S8. Phosphopeptides detected for leghemoglobins

| | | | | Argon | | High nitrate | | nifH | |
|---|---|---|---|---|---|---|---|---|---|
| **AA Sequence** | **Protein ID and Modifications** | **Gene** | **Lb ID** | **q-value** | **fold change** | **q-value** | **fold change** | **q-value** | **fold change** |
| ATGEVVLGDATLGSIHIQK | G7K1Z9 1xPhospho [S92(100)] | MTR5g081030 | Lb2 | 0.00 | **3.87** | 0.00 | **9.17** | 0.00 | **4.07** |
| QEALVNSSWELFK | G7I6B5 1xPhospho [S13(98.3)] | MTR1g011540 | Lb7 | 0.00 | **3.72** | 0.00 | **6.81** | 0.00 | **3.39** |
| QEALVNSSYEAFK | G7KBK3 1xPhospho [S/Y] | MTR5g066070 | Lb9 | 0.00 | **4.23** | 0.00 | **9.85** | 0.00 | **3.26** |
| QEALVNSSYEAFK | G7KBK3 1xPhospho [S14(88.6)] | MTR5g066070 | Lb9 | 0.00 | **4.99** | 0.00 | **9.11** | 0.00 | **3.25** |
| QEALVNSSYEAFK | G7KBK3 1xPhospho [S13(99.7)] | MTR5g066070 | Lb9 | 0.00 | **4.05** | 0.00 | **6.57** | 0.00 | **3.13** |
| DTAGVQDSPK | G7I6B5 1xPhospho [S55(100)] | MTR1g011540 | Lb7 | 0.00 | **3.53** | 0.00 | **3.80** | 0.01 | **2.54** |
| DSAGVQDSPK | G7KGN2 1xPhospho [S56(100)];G7K1Z7 1xPhospho [S55(100)] | MTR5g041610 | Lb6/LB8 | 0.00 | **3.13** | 0.00 | **5.08** | 0.01 | **2.48** |
| QEALVNSSFESFK | G7K1Z7 1xPhospho [S14(90.6)] | MTR5g081000 | Lb8 | 0.00 | **3.66** | 0.00 | **6.85** | 0.01 | **2.27** |
| DSTGVQDSPQLQAHAEK | G7K1Z9 1xPhospho [S/T] | MTR5g081030 | Lb2 | 0.00 | **3.79** | 0.00 | **9.58** | 0.01 | **2.27** |
| DSAGVQDSPQLQAHAEK | G7KBK3 1xPhospho [S50(100)] | MTR5g066070 | Lb9 | 0.01 | **1.75** | 0.00 | **3.57** | 0.02 | **2.14** |
| DSAGVQDSPK | G7KGN2 1xPhospho [S50(100)];G7K1Z7 1xPhospho [S49(100)] | MTR5g041610 | Lb6/LB8 | 0.00 | **2.93** | 0.00 | **2.39** | 0.02 | **2.10** |
| DSAGVQDSPK | G7KGN2 1xPhospho [S56(100)];G7K1Z7 1xPhospho [S55(100)] | MTR5g041610 | Lb6/LB8 | 0.00 | **2.58** | 0.00 | **3.27** | 0.02 | **2.07** |
| QEALVNSSFESFK | G7K1Z7 1xPhospho [S13(100)] | MTR5g081000 | Lb8 | 0.00 | **2.16** | 0.00 | **2.43** | 0.02 | **1.99** |
| DTTGVQDSPQLQAHAAK | G7KGT0 1xPhospho [T51(99.8)] | MTR5g080440 | Lb1 | 0.00 | **2.87** | 0.00 | **6.70** | 0.02 | **1.93** |
| MGFTENQEALVNSSWESFK | G7K1Z9 1xMet-loss [N-Term];1xPhospho [S13(79.1)] | MTR5g081030 | Lb2 | 0.00 | **3.10** | 0.00 | **10.85** | 0.03 | **1.73** |
| DSAGVQDSPK | G7KGN2 1xPhospho [S50(100)];G7K1Z7 1xPhospho [S49(100)] | MTR5g041610 | Lb6/LB8 | 0.00 | **2.34** | 0.15 | **0.61** | 0.05 | **1.55** |
| QEALVNSSFESFK | G7K1Z7 1xPhospho [S17(99.1)] | MTR5g081000 | Lb8 | 0.00 | **3.50** | 0.00 | **7.77** | 0.11 | **1.22** |
| QESLVNSSWESFK | G7KGT0 1xPhospho [S9(100)] | MTR5g080440 | Lb1 | 0.00 | **3.78** | 0.00 | **8.73** | 0.29 | **0.80** |
| QESLVNSSWESFK | G7KGT0 1xPhospho [S17(100)] | MTR5g080440 | Lb1 | 0.00 | **4.35** | 0.00 | **10.06** | 0.35 | **0.71** |
| DTTGVQDSPQLQAHAAK | G7KGT0 1xPhospho [S56(100)] | MTR5g080440 | Lb1 | 0.00 | **2.76** | 0.00 | **6.73** | 0.62 | **0.40** |
| DSAGVQDSPQLQAHAEK | G7KBK3 1xPhospho [S56(100)] | MTR5g066070 | Lb9 | 0.00 | **2.70** | 0.00 | **3.11** | 0.69 | **0.33** |

#### Fig. S7. Alignement of Lb promoters of M. truncatula

| | **Gene model** | **Gene** | **Promoter sequence** |
|---|---|---|---|
| phosphorylated | Medtr5g066070 | LgHb9 | TTTTGTCTC-TTAATAACTACAATGGTCACCTCCACAAGCCATTATATTCTT |
| | Medtr5g081030 | LgHb2 | TCTGGTCTC-TTCATAATTTCAATGGTCATTTCCACAAGCCAATAGATTCTT |
| | Medtr5g081000 | LgHb8 | TATTGTCTCTATAATATTGTCAATAGCCATTTCCACAAGCCAATAGATTCAT |
| | Medtr5g080440 | LgHb1 | TCTTGTCTC-TTAATAATTTTAATGGTCACCGCCACAAGCCAATAGATTCTT |
| | Medtr1g011540 | LgHb7 | TATTGTCTCTTTAATAATGTCAACAGCCATTTCCACAAGCCAATAGATTCTT |
| | Medtr5g041610 | LgHb6 | TCTTGTCTC-TTAATAATGTCAATAGCCACCTCCACAAGCCAACAAATTCTT |
| | Medtr1g090820 | LgHb10 | AATTGTCTC-TTAATATTGCCAATGGCCACCTCTAAATTAAAGACCAATAGA |
| | Medtr1g049330 | LgHb4 | GATTGTCTC-TTAATAATACCAATAGCCATCTCATAATTAAAGGCCAATAAA |
| | Medtr5g080400 | LgHb3 | TCTTGTCTC-TTAATAATTTCAATGGTCACCCCCAGAAGCCAATAGATTCTT |
|  | Medtr1g090810 | LgHb11 | TATTATCTCTT----AATAAAGGTTGCCTCCTCTCAAAGGCCAAGAAATACT |
| | Medtr7g110180 | LgHb12 | TCACGACTC-TTTATAATAGTAATGGACGTTTCCACAAGCCAACAAATTTAA |
| | Medtr5g080900 | LgHb5 | TATTGTTTCTTTGATAATGTCAATGACCATTTTCACGATTCTTTTAAATATA |
| | Consensus dNRE | | **KKYYYY-WTMA** **YAAYRGYCAYYBC** **CCAAYARATT** |

Jiang et al., 2021

#### Fig. S8. Phosphorylation pattern of leghemoglobin under sanctioning conditions

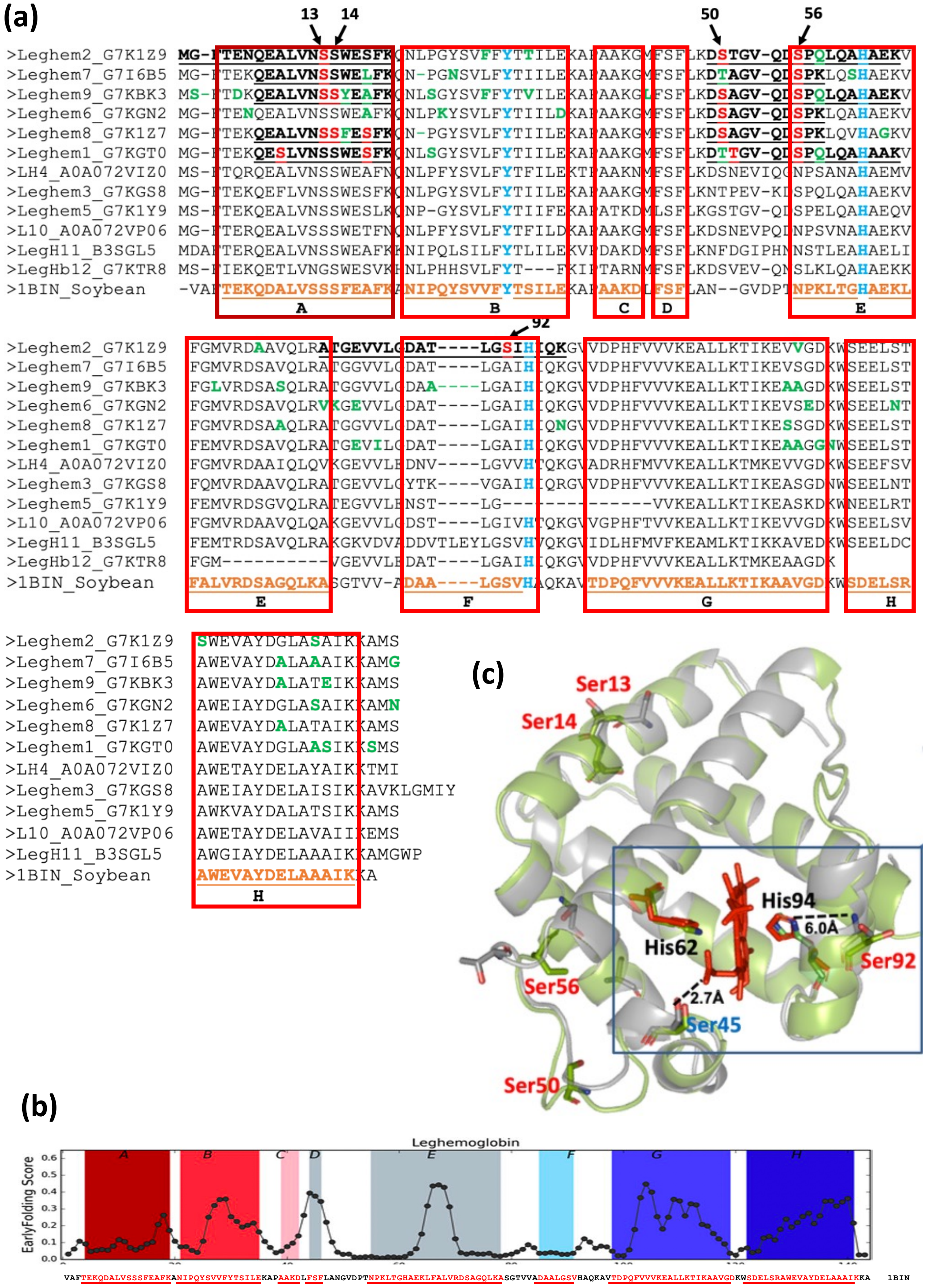

Raimondi, 2017
